# Proteogenomic characterization of hepatocellular carcinoma

**DOI:** 10.1101/2021.03.05.434147

**Authors:** Charlotte K Y Ng, Eva Dazert, Tuyana Boldanova, Mairene Coto-Llerena, Sandro Nuciforo, Caner Ercan, Aleksei Suslov, Marie-Anne Meier, Thomas Bock, Alexander Schmidt, Sylvia Ketterer, Xueya Wang, Stefan Wieland, Matthias S Matter, Marco Colombi, Salvatore Piscuoglio, Luigi M Terracciano, Michael N Hall, Markus H Heim

## Abstract

We performed a proteogenomic analysis of hepatocellular carcinomas (HCCs) across clinical stages and etiologies. We identified pathways differentially regulated on the genomic, transcriptomic, proteomic and phosphoproteomic levels. These pathways are involved in the organization of cellular components, cell cycle control, signaling pathways, transcriptional and translational control and metabolism. Analyses of CNA-mRNA and mRNA-protein correlations identified candidate driver genes involved in epithelial-to-mesenchymal transition, the Wnt-β-catenin pathway, transcriptional control, cholesterol biosynthesis and sphingolipid metabolism. The activity of targetable kinases aurora kinase A and CDKs was upregulated. We found that *CTNNB1* mutations are associated with altered phosphorylation of proteins involved in actin filament organization, whereas *TP53* mutations are associated with elevated CDK1/2/5 activity and altered phosphorylation of proteins involved in lipid and mRNA metabolism. Integrative clustering identified HCC subgroups with distinct regulation of biological processes, metabolic reprogramming and kinase activation. Our analysis provides insights into the molecular processes underlying HCCs.

## INTRODUCTION

Liver cancer was the sixth most commonly diagnosed cancer with 841,000 cases and the fourth leading cause of cancer death with 782,000 deaths globally in 2018 (Arnold et al., 2020). Hepatocellular carcinoma (HCC) accounts for 75%-85% of all primary liver malignancies and has rising incidence and mortality in western countries (Arnold et al., 2020). The past decade has seen numerous studies characterizing the genomic and transcriptomic features and diversity of HCC. Genomic analyses have revealed that *TERT* promoter, *CTNNB1* (encoding β-catenin) and *TP53* (encoding p53) are frequently mutated in HCC, while genes involved in other critical processes, such as oxidative stress response, chromatin remodeling and hepatocyte differentiation, are recurrently mutated but in <10% of HCC (Cancer Genome Atlas Research Network, 2017; Fujimoto et al., 2012, 2016). Transcriptomic subtyping has revealed between 2 and 6 HCC subclasses that differ in the expression of genes related to proliferation, stemness, metabolism, hepatocyte differentiation and liver function (Bidkhori et al., 2018; Boyault et al., 2007; Désert et al., 2017; Hoshida et al., 2009; Lee et al., 2004; Makowska et al., 2016).

More recently, global proteome and phosphoproteome profiling has been made possible by mass spectrometry-based methods. Two proteogenomic studies of HCCs, both of hepatitis B virus (HBV)-associated HCCs, have been published (Gao et al., 2019; Jiang et al., 2019). In the first study, the proteome and phosphoproteome profiling of early-stage HBV-associated HCCs found that a subset of HCCs characterized by disrupted cholesterol homeostasis and overexpression of SOAT1 was associated with poor outcome (Jiang et al., 2019). Indeed, avasimibe, a SOAT1 inhibitor, effectively reduced the size of tumors that overexpressed SOAT1 in patient-derived xenograft mouse models (Jiang et al., 2019). In the second study, integrated proteogenomic analysis of HBV-related HCC (Gao et al., 2019) revealed activation status of key signaling pathways and metabolic reprogramming in HBV-related HCC. In particular, the authors identified three proteome subclasses, namely metabolism, proliferation and microenvironment dysregulated subgroups, that were associated with clinical and molecular features such as patient survival, tumor thrombus and genetic profile. PYCR2 and ADH1A, both involved in metabolic reprogramming in HCC, were further identified as proteomic prognostic biomarkers. In contrast to the surgically resected HBV-associated HCCs profiled in the previous studies, here we performed an integrated proteogenomic analysis of HCC biopsies across diverse etiologies and clinical stages.

## RESULTS

### Proteogenomic profiling of HCC

We collected paired biopsies from 122 tumors and 115 non-tumoral tissues from 114 patients (Table 1 **and Table S1**). Six patients had 2 synchronous multicentric tumor biopsies with a single matched non-tumoral tissue included in the study. One patient had 3 multicentric tumor biopsies and two non-tumoral biopsies, obtained 7 years apart, included in the study (**Figure S1**). None of the patients had undergone systemic therapy for their disease. At the time of the study, 53% of the patients were early stage (BCLC 0/A) and 47% were advanced stage (BCLC B/C/D). 94% of the patients had at least one underlying liver disease, with alcohol liver disease (59%) and hepatitis C infection (26%) being the most common. Most biopsies were of intermediate grade (54% Edmondson grade 2 and 34% grade 3). Whole-exome sequencing and RNA-sequencing was performed for all 122 tumors and 115 non-tumoral biopsies (**Table S1**). A subset of 51 tumors from 49 patients were subjected to global proteome and phosphoproteome profiling using liquid chromatography-tandem MS analyses (Table 1 **and Table S1**). The proteome was measured in data-independent manner by selected window acquisition of theoretical mass (SWATH) (Gillet et al., 2012) adapted for HCC biopsies (Guri et al., 2017), which is an efficient method to acquire quantitative MS data of a large cohort in a relatively short time frame. The phosphoproteome was measured in data-dependent and label-free manner. Aside from a slight enrichment of HCV-associated etiology (p=0.04), no other difference in terms of the clinicopathological parameters was observed in the 51 tumors subjected to proteome and phosphoproteome profiling compared to the 71 that were not (all p>0.05, Table 1) and no difference in terms of their mutational and transcriptomic landscape (**Figure S2**). As controls, we also performed RNA-sequencing, global proteome and phosphoproteome profiling on 15, 11 and 10 normal biopsies from 19 patients without HCC and with normal liver values, respectively (**Table S1**).

**Table 1:**
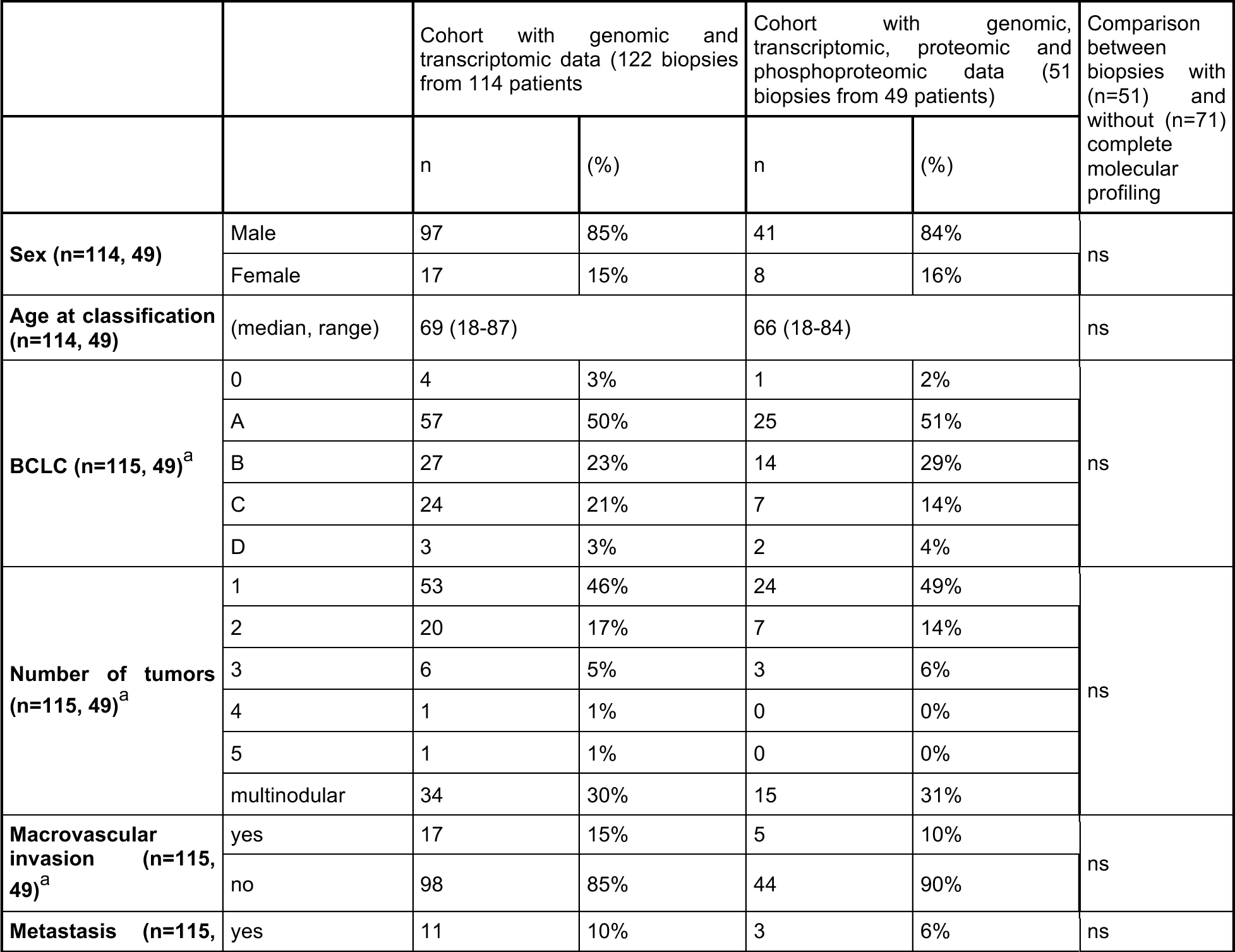
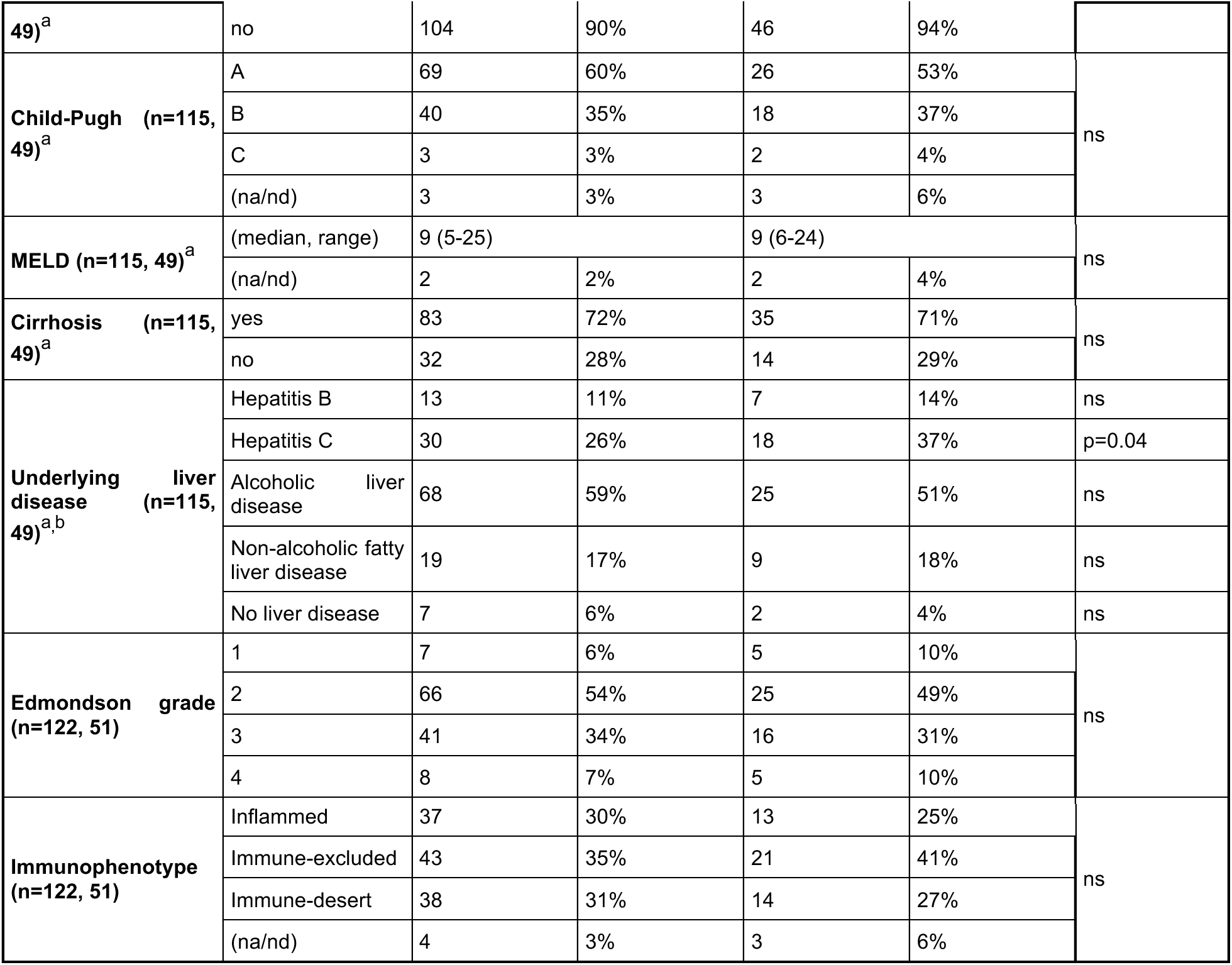
Summary of clinicopathological information of the cohort. ^a^Determined at the time of biopsy. One patient was biopsied twice seven years apart. ^b^Patient may have >1 underlying liver disease. Statistical comparisons were performed using Fisher’s exact tests (for categorical data with two levels), Chi-squared tests (for categorical data with >2 levels), and Mann-Whitney U tests (for numerical and ordinal data). BCLC: Barcelona Clinic Liver Cancer clinical staging system; MELD: model for end-stage liver disease; na: not available; nd: not determined; ns: not significant. See also Figures S1-2 and Table S1.

### Deregulated pathways in HCC

Principal component analyses showed that HCCs are distinct from and more variable than normal livers on the transcriptome, proteome and phosphoproteome levels, with the normal livers forming a tight cluster and the HCCs spread out (Figure 1A-C). Across all molecular levels, low-grade HCCs were more homogeneous than high-grade HCCs as measured by intra-group variability (Spearman rho between 0.26 and 0.37, all p<0.0001, Figure 1D). In accordance with the definition of histological (Edmondson) grading, we also found that low-grade HCCs were more similar to normal livers than high-grade HCCs (Spearman rho between 0.51 and 0.66, all p<0.0001, Figure 1E).

**Figure 1:**
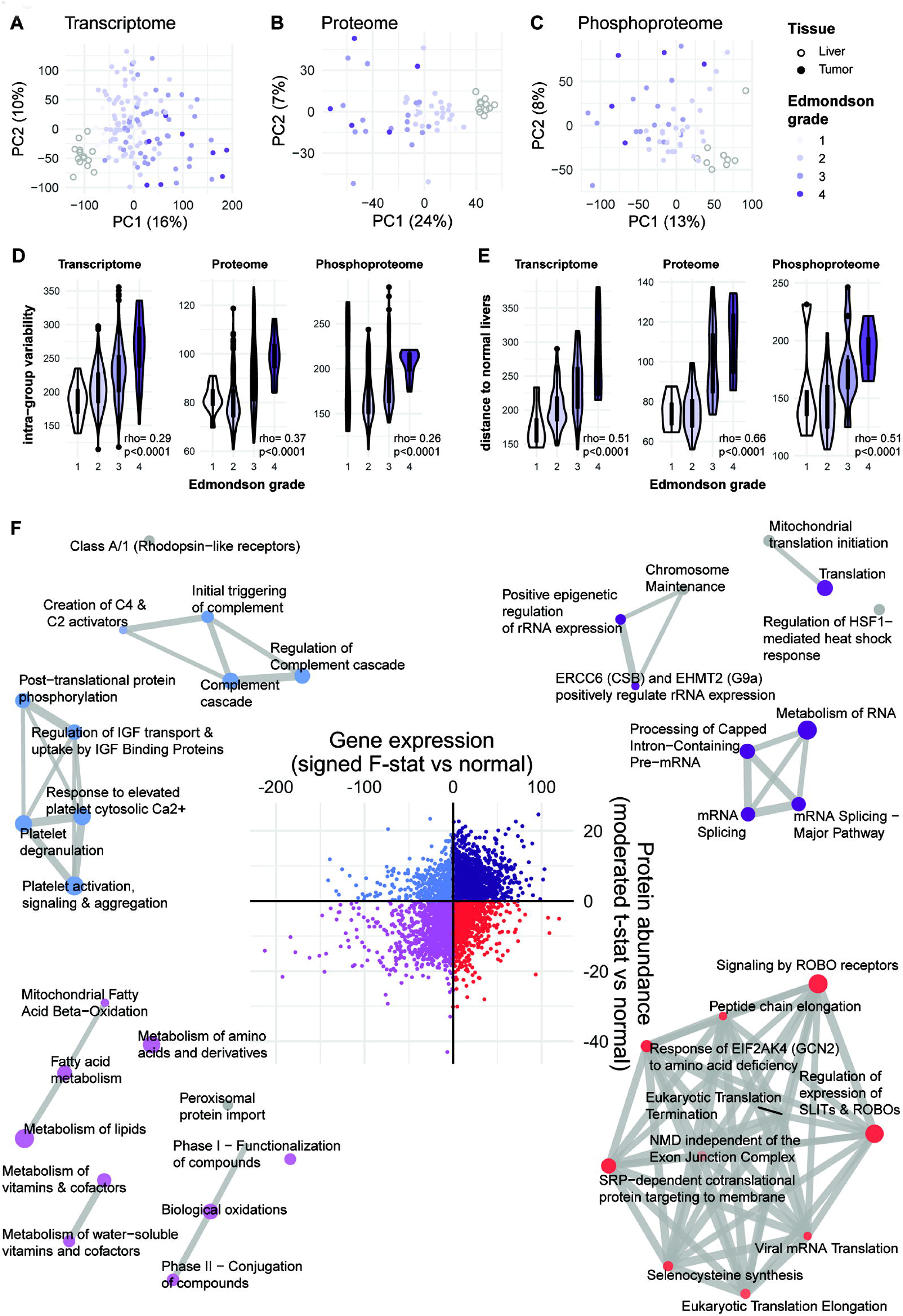
Deregulated pathways in HCC. **(A-C)** Principal component analysis plots of (**A**) transcriptome, (**B**) proteome, (**C**) phosphoproteome of HCC biopsies (colored by Edmondson grade) and normal liver biopsies. **(D)** Intra-group (within Edmondson grade) variability as measured by pairwise Euclidean distance between samples according to principal components. **(E)** Distance of each HCC to the median of normal livers as measured by Euclidean distance according to principal components. (**F**) Scatter plot of (y-axis) the moderated t-statistics from the differential protein expression analysis of HCC vs normal liver against (x-axis) the F-statistics from the differential gene expression analysis of HCC vs normal liver. Points are colored according to the four quadrants. Enrichment maps show the top 10 enriched Reactome pathways from over-representation tests of the genes/proteins in each of the four quadrants. In each enrichment map, gene sets with overlapping gene sets are joined by edges. Nodes are colored according to p-value, where gray indicates a higher p-value and dark blue/violet/purple/red indicates a lower p-value. The size of the nodes is proportional to the number of genes in the quadrant within a given gene set. See also Table S2.

To identify pathways deregulated in HCC, we performed differential expression analyses of the transcriptome and proteome of HCCs compared to normal livers. Overall, we observed a moderate correlation between the deregulation of the transcriptome and the proteome (Spearman rho=0.33, p<0.0001, Figure 1F). We performed a quadrant analysis of transcriptome and proteome data to identify genes and proteins consistently up-/down-regulated, or regulated differently. We found that 37.7% (15.7% for adjusted p≤0.05) of genes were up-regulated on both levels, 20.9% (6.2%) were up-regulated on the mRNA level but down-regulated on the protein level, 16.6% (5.4%) were up-regulated in protein but down-regulated on the mRNA level, and 24.8% (13.6%) were down-regulated in both. Pathway analysis of these four quadrants revealed that the genes that were up-regulated on both the mRNA and the protein levels were enriched in pathways related to mRNA splicing, epigenetic regulation of rRNA expression and translation (Figure 1F **and Table S2**). By contrast, genes that were consistently down-regulated were enriched in pathways related to metabolism of amino acids, fatty acids, xenobiotics and other metabolites. Among the genes up-regulated only on the mRNA level, pathways related to translational control, proteasome and oxidative phosphorylation were enriched. By contrast, pathways related to the complement and coagulation were enriched among genes upregulated only on the protein level.

Taken together, while we observed overall upregulation of pathways related to mRNA splicing and downregulation of pathways related to normal liver function, we also observed pathways related to translational control being upregulated on the mRNA level only, and pathways related to coagulation and the complement upregulated on the protein level only.

### CNA-mRNA-protein correlation analysis identifies candidate driver genes

Next, we evaluated the correlation between copy number alteration (CNA), mRNA expression and protein expression. The median CNA-mRNA and mRNA-protein Spearman correlation coefficients were 0.201 and 0.282, respectively (Figure 2A-B). For the CNA-mRNA correlation, 50.10% of the genes showed significant positive correlation, while for the mRNA-protein correlation, 45.08% of the genes showed significant positive correlation. Gene set enrichment analysis of CNA-mRNA and mRNA-protein expression correlation revealed 275 and 45 overlapping Reactome pathways, with only one (protein localization) enriched in both comparisons. The pathways enriched among genes that showed high CNA-mRNA correlation include RNA transport, ubiquitination and proteasome degradation, transcriptional regulation by *TP53*, translation, cell cycle and DNA repair, cellular response to stress and asparagine N-linked glycosylation (Figure 2C **and Table S3**). By contrast, genes that showed high mRNA-protein expression correlation are enriched in pathways related to the metabolism of amino acids, glucose, fatty acids, xenobiotics and biological oxidations (Figure 2C **and Table S3**).

**Figure 2:**
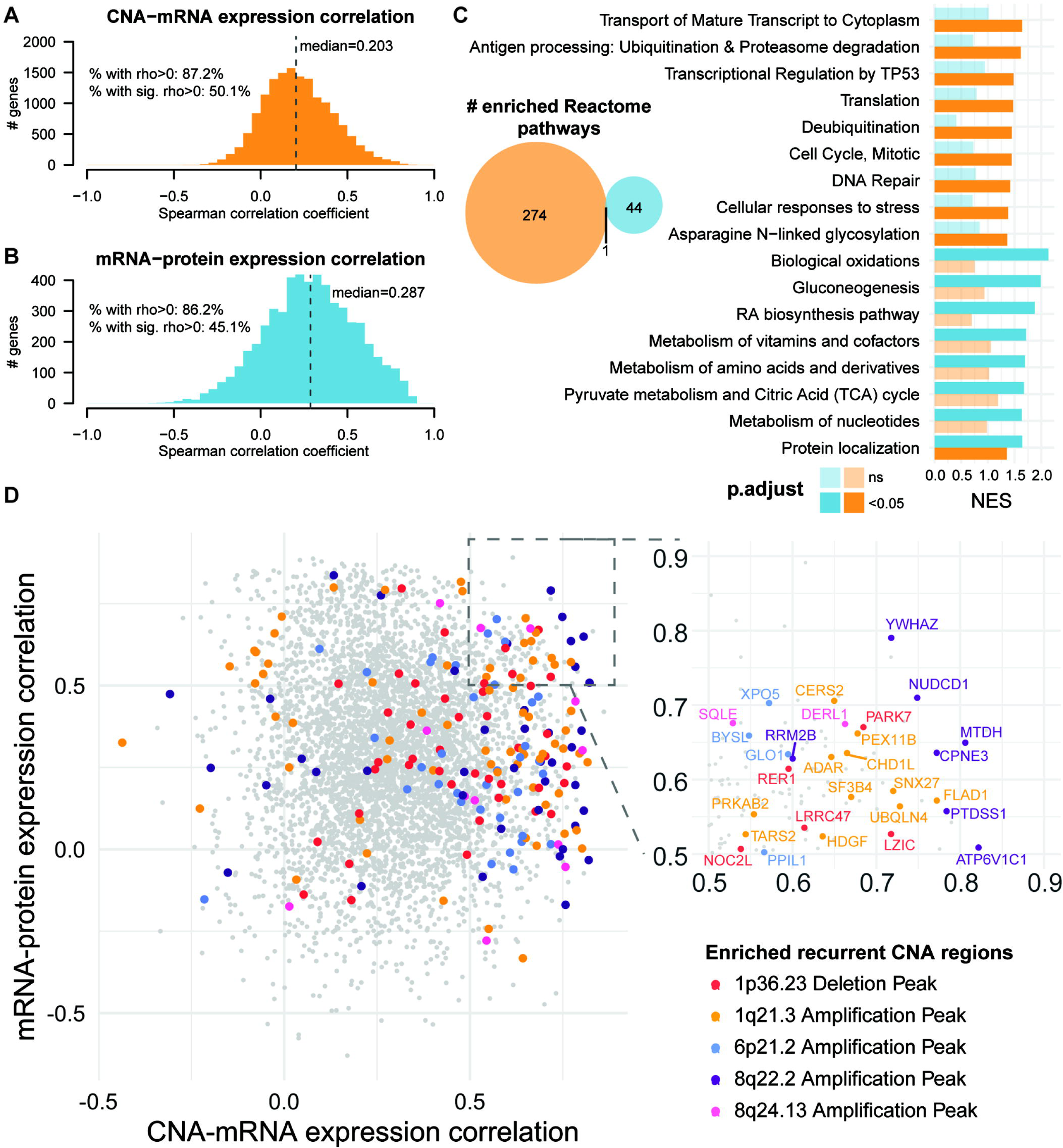
CNA-mRNA-protein correlation. **(A-B)** Histograms of the distributions of the per-gene Spearman correlation coefficients (**A**) for CNA-mRNA and (**B**) mRNA-protein expression. sig.: significant. (**C**) Venn diagram of the number of enriched Reactome pathways for genes/proteins ranked by CNA-mRNA expression correlation (orange) and mRNA-protein expression correlation (blue). Enrichment was defined by gene set enrichment analysis. Barplot of selected Reactome pathways enriched among genes with high CNA-mRNA expression correlation (orange) and/or with high mRNA-protein expression correlation (blue). Statistically significant normalized enrichment scores (NES, p<0.05) are shown in darker shades (dark orange/blue) while non-significant NESs are shown in lighter shades (light orange/blue). (**D**) Scatterplot of the per-gene Spearman correlation coefficients (y-axis) between mRNA and protein expression against (x-axis) between CNA and mRNA. Genes in five of the recurrently altered regions as defined by GISTIC2 are colored according to the color key. Inset shows the genes with >0.5 correlation coefficients in both comparisons. See also Tables S3-4.

Genome-wide copy number analysis identified 8 recurrently amplified peaks and 11 recurrently deleted peaks (**Table S3**), 5 of which were enriched among genes with high CNA-mRNA correlation but none was enriched among genes with high mRNA-protein correlation (Figure 2D **and Table S3**). While many of the genes in these recurrently amplified or deleted genomic regions showed high correlation between CNA and mRNA, many of them showed little-to-no mRNA-protein correlation (Figure 2D). One could hypothesize that candidate driver genes would be over-represented among *cis*-copy number-regulated genes (i.e. when the copy number variant impacts its own expression) that also show high mRNA-protein correlation. To identify such candidate driver genes within the recurrently altered regions, we focused on the 136 genes that showed high CNA-mRNA and mRNA-protein correlation (Spearman rho>0.5, **Table S4**). Among this group of genes were known cancer genes and putative drug targets such as *CHD1L* (Chromodomain Helicase DNA Binding Protein 1 Like, 1q21.3 peak) (Cheng et al., 2013), *ADAR* (Adenosine Deaminase RNA Specific, 1q21.3 peak) (Chan et al., 2014), *MTDH* (metadherin, 8q22.2 peak) (Dhiman et al., 2019; Hu et al., 2009; Shi and Wang, 2015), and *YWHAZ* (also known as 14-3-3 zeta, 8q22.2 peak) (Pennington et al., 2018; Yang et al., 2012) (Figure 2D **inset**).

There were other candidate copy number-driven cancer genes implicated in oncogenesis (Figure 2D **inset and Table S4**). For example, in addition to *YWHAZ*, the 8q22.2 amplicon also contains *NUDCD1* (NudC domain containing 1, also known as OVA66), previously shown to promote colorectal carcinogenesis and metastasis by inducing EMT and inhibiting apoptosis (Han et al., 2018) and to promote oncogenic transformation by hyperactivating the PI3K/AKT, ERK1/2-MAPK and IGF-1R-MAPK signaling pathways (Rao et al., 2014a, 2014b). The overexpression of *SQLE* (Squalene epoxidase, 8q24.13 peak), an enzyme involved in cholesterol biosynthesis, promotes cell proliferation and migration in HCC cells and positively regulates ERK signaling (Sui et al., 2015). *UBQLN4* (Ubiquilin-4, 1q21.3 peak) has recently been suggested to regulate Wnt-β-catenin pathway activation in HCC cells (Yu et al., 2020) and is associated with genomic instability (Jachimowicz et al., 2019). Of note, there are many poorly characterized genes among those that showed strong CNA-mRNA and mRNA-protein correlation, such as *ALYREF*, involved in transcriptional control and mRNA stabilization (Hautbergue et al., 2008; Hung et al., 2010; Stubbs and Conrad, 2015; Stubbs et al., 2012) and whose expression correlated with cell cycle regulation and mitosis and poor prognosis in HCC (He et al., 2020), and *CERS2*, a key component in sphingolipid metabolism, whose co-expression with TGF-β1 is associated with poor outcome in HCC (Ruan et al., 2016). The role of these genes/proteins in HCC has not been studied in depth but may warrant further investigation. On the other hand, *MTOR* in the 1p36.23 deletion peak showed high CNA-mRNA correlation (rho=0.58) but no mRNA-protein correlation (p>0.05), suggesting alternative mechanisms (e.g. epigenetic) for the regulation of protein expression.

Taken together, our analysis of the CNA-mRNA and mRNA-protein expression correlations showed distinct pathways being regulated on different levels and identified potential HCC driver genes.

### Dysregulated phosphorylation in HCC

Next, we investigated the protein phosphorylation landscape in HCC. Given that protein phosphorylation may be highly driven by the protein expression rather than changes in phosphorylation, we investigated dysregulated phosphorylation sites with and without normalization by overall protein levels. Differential expression analyses revealed 692 and 648 sites that are hyper- and hypophosphorylated, respectively, and 302 and 355 normalized (by overall protein levels) hyper- and hypophosphorylated sites compared to normal livers (adjusted p≤0.05 and |log_2_ fold-change|>1, Figure 3A**, Figure S3A and Table S5**). A pathway enrichment analysis revealed that the hyperphosphorylated sites are in proteins involved in cell cycle, mRNA splicing, the immune system, cancer-related signaling pathways such as receptor tyrosine kinases and MAP kinase and regulation by PTEN and p53 (Figure 3B**, Figure S3B and Table S6**). Signaling by AKT, FGFR, VEGF, TGF-beta are also enriched among the hyperphosphorylated proteins, though not always statistically significant in the analyses of both normalized and unnormalized phosphorylation levels. Among the hypophosphorylated sites, processes related to histone modification were enriched (Figure 3B**, Figure S3B and Table S6**). Interestingly, pathways related to metabolism of amino acids, carbohydrates, lipids were enriched among the overall hypophosphorylated sites but also the normalized hyperphosphorylated sites. By contrast, proteins involved cellular senescence and chromatin organization showed the opposite trend, with enrichment of overall hyperphosphorylated sites but normalized hypophosphorylated sites. While base excision repair and DNA double-strand break repair were enriched among both hypophosphorylated sites, nucleotide excision repair was enriched among the hyperphosphorylated sites (Figure 3B**, Figure S3B and Table S6**).

**Figure 3:**
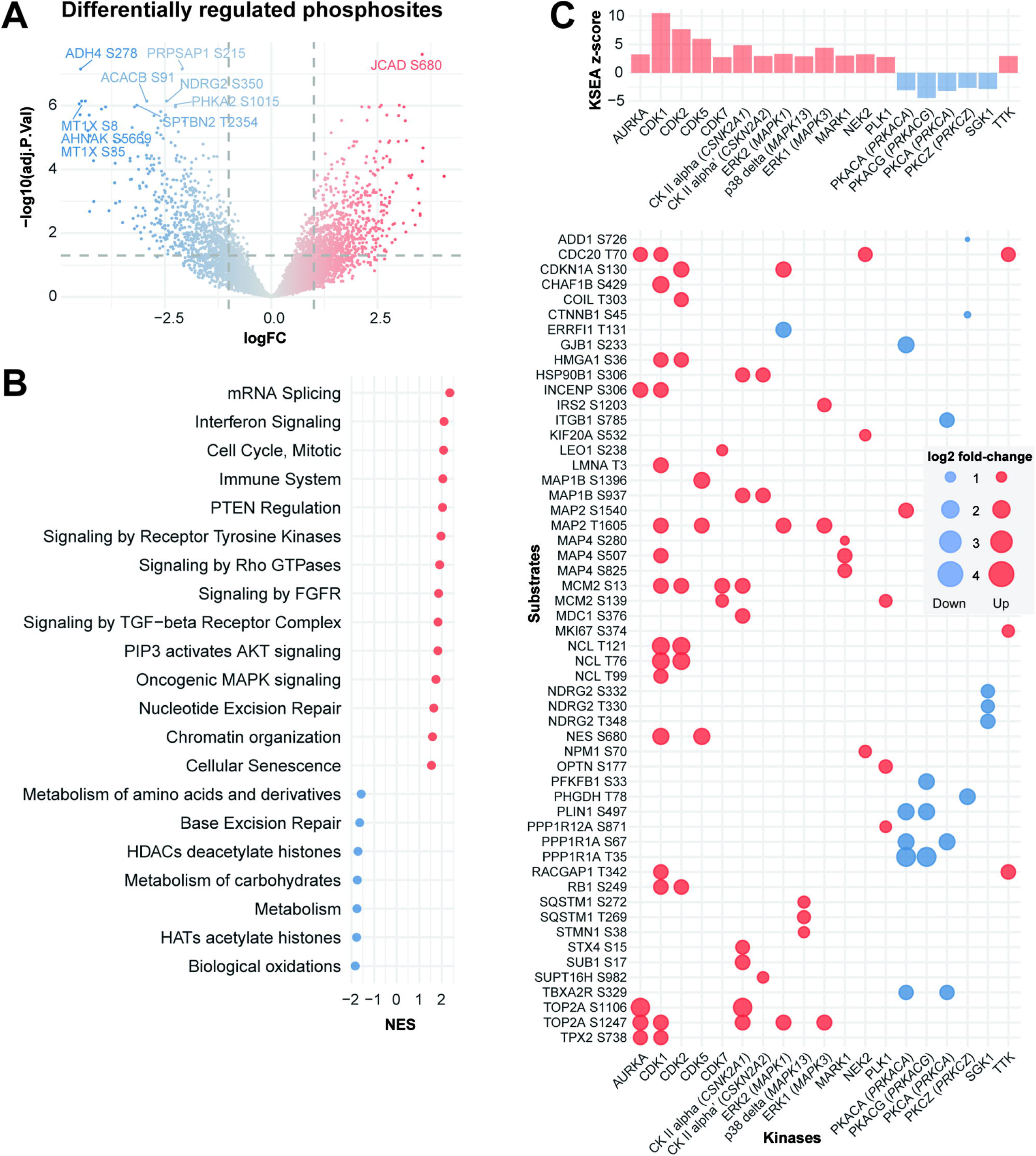
The phosphoproteomic landscape of HCC. **(A)** Volcano plot of the −log10(adjusted p-value) against the log fold-change (logFC) of the differentially regulated phosphorylation sites in HCC compared to normal livers. Dots are colored by logFC. Vertical dotted lines indicate |logFC|=2 and horizontal dotted lines indicate adjusted p-value=0.05. **(B)** Dot plot illustrating selected enriched Reactome pathways according to gene set enrichment analysis (GSEA) from the differential expression analysis in (**A**). NES: normalized enrichment score. **(C)** Top barplot showing the enrichment z-score of the kinases with significantly up- or downregulated kinase activity in a kinase-substrate enrichment analysis (KSEA) comparing HCC to normal livers. In the bubble plot below, the phosphorylation site substrates are shown in rows, where red and blue dots indicate that the phosphorylation site is up- and downregulated, respectively. The size of the dots is proportional to the log_2_ fold-change of the phosphorylation site. Phosphorylation sites with at least a 5-fold difference between HCCs and normal livers are shown. For kinases with <3 substrates with at least a 5-fold difference, the top three substrates with the highest |logFC| are shown. See also Figure S3 and Tables S5-7.

To infer the activation of kinases in HCC, we performed a Kinase-Substrate Enrichment Analysis (KSEA)(Casado et al., 2013). KSEA revealed that Aurora kinase A (*AURKA*), Cyclin-dependent kinases 1/2/5/7 (*CDK1/2/5/7*), ERK1/2 (*MAPK1/3*) and PLK1 showed increased activation compared to normal livers, while PKACA/G (*PRKACA/G*), PKCA/Z (PRKCA/Z) and SGK1 showed reduced activity (Figure 3C **and Table S7**). When analyzing dysregulated phosphorylation normalized by protein level, KSEA revealed increased AURKA, CDK1/2/5 and ERK1/2 but also GSK3B activity in HCCs (**Figure S3C and Table S7**).

Our results show that altered phosphorylation in HCC affects a wide range of biological processes from cell proliferation and DNA repair to immune system and signal transduction pathways.

### Proteogenomic analysis of significantly mutated genes

Using whole-exome sequencing, we identified 24,488 somatic mutations (23,660 SNVs and 828 indels) across the 122 tumor biopsies (**Figure S4A and Table S8**). One tumor (D096) was hypermutated with 9035 mutations. In the remaining 121 tumor biopsies, we identified a median of 123 somatic mutations per biopsy (range 25-446). Using MutSigCV (Lawrence et al., 2013) and OncodriveFML (Mularoni et al., 2016), we identified 7 significantly mutated genes (SMGs, *ALB*, *ARID1A*, *AXIN1, CDKN1A*, *CTNNB1*, *GPAM*, *TP53*, **Figure S4A**). While *ALB*, *ARID1A*, *AXIN1, CDKN1A*, *CTNNB1* and *TP53* had previously been identified as SMGs in more than one genomic study (Bailey et al., 2018; Fujimoto et al., 2016; Martincorena et al., 2018; Schulze et al., 2015), *GPAM* was identified as an SMG only in a meta-analysis of HCC genomic studies (Li et al., 2018) (1.8% vs 7.4% in the current study, Fisher’s exact test, p=0.001). Here we found seven of the nine *GPAM* mutations were frameshift mutations, strongly suggestive of a tumor suppressor role (**Figure S4B**), though only one was homozygous (bi-allelic inactivation).

We evaluated the clinicopathological correlates of the 7 SMGs, together with 7 additional cancer genes identified from at least 2 previous HCC genomics studies (Bailey et al., 2018; Fujimoto et al., 2016; Martincorena et al., 2018; Schulze et al., 2015) and mutated in at least 3 HCCs of the current cohort (*ACVR2A*, *APOB*, *ARID2*, *CDKN2A*, *KEAP1*, *RB1* and *TSC2*). We found that *CTNNB1*-mutant tumors were more frequently lower grade (Edmondson grade, p=0.017) and with a pseudoglandular growth pattern (p=0.034). Consistent with previous reports (Calderaro et al., 2017; Luke et al., 2019), *CTNNB1*-mutant tumors were also associated with the immune-desert phenotype (p=0.039). By contrast, *TP53*-mutant tumors were of higher grade (p=0.001) and associated with HBV (p=0.010, **Figure S4C**). However, none of the clinicopathological associations was statistically significant after correcting for multiple testing. Among the 14 driver genes, *TP53* and *CTNNB1* mutations were mutually exclusive (odds ratio=0.35, p=0.012, Fisher’s exact test), as previously reported (Guichard et al., 2012). A multivariate Cox-proportional hazard model suggests that mutations in *CDKN2A*, *GPAM*, *KEAP1* and *TSC2* are associated with poor overall survival independent of BCLC clinical stage (**Figure S4D**).

To evaluate the transcriptome, proteome and phosphoproteome changes associated with *CTNNB1* and *TP53* mutations, we performed differential expression analyses comparing mutant and wild-type HCCs. We identified 3067 differentially expressed genes in *CTNNB1*-mutant and 3949 in *TP53*-mutant HCCs. Changes on the protein level were also observed, with 23 differentially expressed proteins for *CTNNB1*-mutant and 399 for *TP53*-mutant HCCs (Figure 4A,E and **Table S9**). No statistically significant differences were observed on the phosphoproteome level at FDR 5%. Of the 23 proteins that were up- or down-regulated in *CTNNB1*-mutant HCCs, 13 were also differentially expressed at the mRNA level (Figure 4A and **Table S9**). These include glutamine synthetase (encoded by *GLUL*), α-methylacyl-CoA racemase (*AMACR*, reported to be associated with *CTNNB1* mutations in HCC (Sekine et al., 2011)), ACSS3 (*ACSS3*, associated with a metabolic HCC subclass characterized by frequent *CTNNB1* mutations (Bidkhori et al., 2018)). On the other hand, the remaining ten differentially expressed proteins were not associated with differential transcription. Notably, these include TNRC6B (*TNRC6B*, involved in the β-catenin-independent Wnt signaling), Protein Kinase C Epsilon (*PRKCE*, a β-catenin binding partner (Duong et al., 2017)), and PPIE (*PPIE*, a spliceosome component that regulates the splicing of the long non-coding RNA *FAST* which in turns regulates β-catenin and Wnt signaling (Guo et al., 2020)). Several of the Wnt target genes whose mRNA expression is typically altered in association with *CTNNB1* mutation, such as *NKD1*, *AXIN2, RNF43* and *ALDH3A1* are not differentially expressed at the protein level (Figure 4A and **Table S9**).

**Figure 4:**
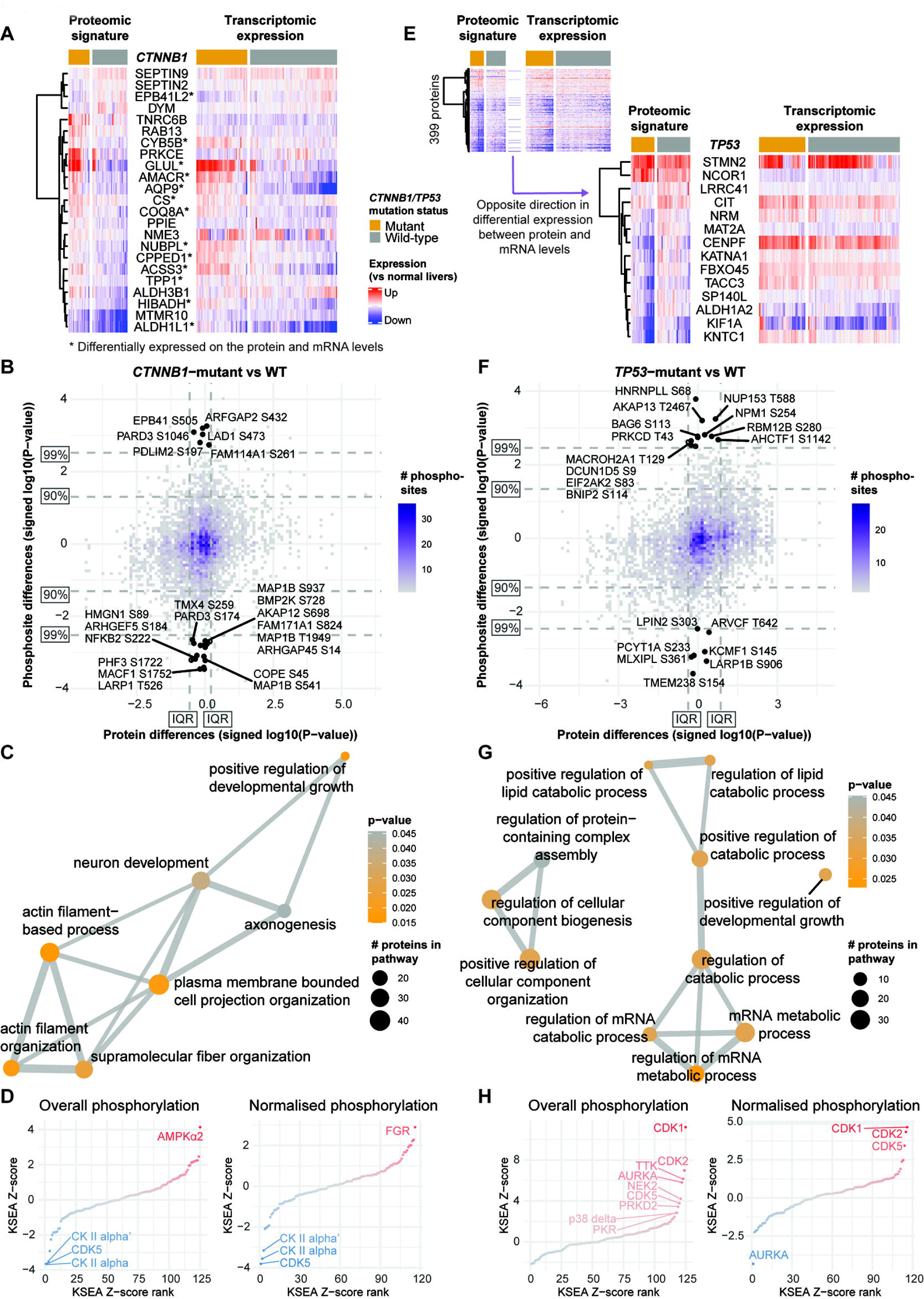
Proteogenomic analysis of SMGs. **(A)** Heatmaps showing the log fold-change (compared to the median of normal livers) of (left) 23 proteins differentially expressed between *CTNNB1*-mutant and -WT HCCs (FDR<0.05) and (right) the corresponding gene expression on the mRNA level. Samples are stratified according to *CTNNB1* mutation status. Genes with asterisks were also differentially expressed on the transcriptome level. (**B**) Binned scatterplot plotting the signed p-values from differential expression analyses of protein expression (x-axis) and of phosphorylation site expression (y-axis) between *CTNNB1*-mutant and -WT HCCs. Signed p-values refer to the p-values from differential expression analyses signed according to the direction of the fold change. Phosphorylation sites at >99th quantile of the unsigned p-values with their corresponding protein within the inter-quartile range of signed p-values of differential protein expression analysis are labeled. **(C)** Enrichment map showing the Gene Ontology biological processes enriched among proteins with phosphorylation sites at >90th quantile of the unsigned p-values with their corresponding protein within the inter-quartile range of signed p-values of differential protein expression analysis. **(D)** Plot showing the kinase-substrate enrichment analyses (KSEA) enrichment z-scores ordered in increasing order, comparing (left) phosphorylation site abundance and (right) phosphorylation site abundance normalized by protein abundance between *CTNNB1*-mutant vs -WT HCCs. Significant kinases are labeled. **(E)** Small heatmap showing the log fold-change (compared to the median of normal livers) of (left) 399 proteins differentially expressed between *TP53*-mutant and -WT HCCs (FDR<0.05) and (right) the corresponding gene expression on the mRNA level. Large heatmap showing the subset of 14 proteins/genes for which the direction of the differential expression between *TP53*-mutant and - WT HCCs differed between the proteomic and transcriptomic signatures. Samples are stratified according to *TP53* mutation status. **(F-H)** as **(B-D)** for stratified by *TP53* mutation status. See also Figure S4 and Tables S8-10.

*CTNNB1* encodes β-catenin, a protein involved in intercellular adhesion. In HCC, mutations in *CTNNB1* lead to accumulation of cytoplasmic β-catenin and its subsequent relocation to the nucleus and aberrant activation of the Wnt pathway. Therefore, we searched for phosphorylations that differ the most between *CTNNB1*-mutant and *CTNNB1*-wild-type HCCs (10% in terms of p-value from differential expression analysis) but are not associated with differences on the protein level (interquartile range in terms of p-value signed according to the direction of differential expression, Figure 4B). Pathway analysis showed that 189 such phosphorylation sites are in proteins involved in the regulation of actin filament organization and related processes (Figure 4C **and Table S10**). Among the sites that showed increased phosphorylation in *CTNNB1*-mutant HCCs were Par3-alpha (encoded by *PARD3*) S1046 and PDLIM2 (*PDLIM2*) S197. On the other hand, MAP1B (*MAP1B*) S541/S937/S1396 and ACF7 (*MACF1*) S1752 showed decreased phosphorylation in *CTNNB1*-mutant HCCs. Phosphorylation or loss of Par3-alpha has been found to lead to the loss of cell polarity (McCaffrey et al., 2016; Xue et al., 2013). Similarly, phosphorylation of PDLIM2 leads to its stabilization in the cytoplasm and facilitates β-catenin activation (Cox et al., 2019). By contrast, Wnt activation causes GSK3 kinase inactivation and decreased MAP1B and ACF7 phosphorylation, resulting in increased microtubule stability (Trivedi et al., 2005) and migration (Wu et al., 2011; Zaoui et al., 2010). KSEA revealed that, compared to *CTNNB1-*wild-type HCCs, *CTNNB1*-mutant HCCs showed increased kinase activity of AMPKα2 and reduced activity of the kinases CK II alpha, CK II alpha’ and CDK5 (Figure 4D). On the other hand, KSEA of the normalized phosphorylation levels showed increased activity of FGR, a kinase that contributes to the regulation of immune response and cytoskeleton remodeling (Figure 4D).

Similarly, of the 399 differentially expressed proteins between *TP53*-mutant and -wild-type HCCs, 238 were also significantly differentially expressed at the mRNA level (**Table S9**). Interestingly, the direction of the protein differential expression for 14 of these genes/proteins differed on the mRNA level. While stathmin 2 (*STMN2*) and Nuclear receptor corepressor 1 (*NCOR1*) were overexpressed at the protein level, they were underexpressed on the mRNA level (Figure 4E). By contrast, the remaining 12 were underexpressed on the protein level but overexpressed on the mRNA level, and these included Centromere Protein F (*CENPF*), TACC3 (*TACC3*) and Kinetochore-associated protein 1 (*KNTC1*), all involved in the regulation of the mitotic spindle. Focusing on sites whose phosphorylation differed between *TP53*-mutant and -wild-type HCCs without differences on the protein level, we identified 178 such phosphorylation sites (Figure 4F). A pathway analysis of these phosphorylation sites suggests that *TP53* mutations are associated with phosphorylation changes in proteins involved in the regulation of lipid and mRNA metabolic processes, and the regulation of cellular component biogenesis and organization (Figure 4G **and Table S10**). In particular, PKR (*EIF2AK2*) S83 is an activating autophosphorylation site (Taylor et al., 2001). PKR is involved in diverse cellular processes, including stress response against pathogens (e.g., HCV) and its activation inhibits protein synthesis (García et al., 2006). HCV infection triggers PKR phosphorylation (Garaigorta and Chisari, 2009), though here we did not observe an enrichment of HCV-associated HCCs among the *TP53*-mutant HCCs (p>0.05, Fisher’s exact test). KSEA revealed that *TP53*-mutant HCCs showed increased activity of the kinases CDK1/2/5 (Figure 4H). Several other kinases involved in the control of cell cycle, mitotic checkpoint and spindle formation (Aurora Kinase A, TTK, NEK2), protein synthesis and stress response (PKR), and MAPK signaling (PRKD2, p38 delta (*MAPK13*)) were also found to show increased activity when comparing overall phosphorylation levels. Interestingly, when comparing the normalized phosphorylation levels between *TP53*-mutant and -wild-type HCCs, Aurora Kinase A was found to show reduced pathway activation.

Taken together, the proteogenomic analysis of *CTNNB1* mutations in HCC suggest that the EMT phenotype frequently seen in *CTNNB1*-mutant HCCs may result from alterations in phosphorylation in proteins involved in pathways related to the organization of actin filaments, thereby regulating cell polarity and migration. On the other hand, *TP53*-mutant HCCs are associated with altered phosphorylation of proteins related to cell cycle control, spindle formation and protein synthesis.

### Molecular subtypes of HCC

While molecular subtyping has been performed for transcriptome and proteome data, integrative clustering incorporating phosphoproteome data has not been performed. We therefore performed unsupervised analyses to identify HCC subtypes in each of the 5 individual omics data sets (‘single-omics’) as well as an integrative analysis incorporating all 5 data sets, namely, somatic mutation, CNA, transcriptome, proteome and phosphoproteome (n=122 for the first three types and n=51 for the last two, **Figure S5A-E**). For the single-omics analyses, we identified between 2 and 4 robust clusters using two independent clustering approaches for each of the 5 omics types. On the somatic mutation level, we identified four subclasses characterized by the presence of a mutation in *CTNNB1*, *TP53*, or *ARID1A* or the lack of a mutation in these three genes (**Figure S5F**), while on the CNA level, two subclasses were identified, distinguished by the overall level of genomic instability (**Figure S5G**). For the transcriptome, we identified three subclasses that were distinguished by elevated cell cycle related processes, immune pathways or metabolism (**Figure S5H and Table S11**). Of note, the mutation and transcriptome clusters were associated with Edmondson grade (both p<0.05, chi-squared tests).

For the proteome and phosphoproteome, in each case two clusters were identified. The first proteome subclass is associated with ribonucleoprotein organization, as well as mRNA splicing and processing, while the second subclass is associated with metabolism, peroxisome organization and oxidative phosphorylation (Figure 5A **and Table S11**). The first phosphoproteome subclass is also linked to ribonucleoprotein assembly and mRNA splicing and processing but also to chromatin organization and activation of CK II alpha. On the other hand, the second phosphoproteome subclass is associated with processes such as telomere maintenance, nucleosome organization, DNA repair, actin cytoskeleton regulation and activation of PKA C-alpha (Figure 5B **and Table S11**). While the proteome clusters are associated with Edmondson grade (p<0.05, Fisher’s exact test), the phosphoproteome clusters are not (p>0.05, Fisher’s exact test). A comparison between the various single-omics clusterings revealed significant association between mutation, transcriptome and proteome clusterings (i.e. those that were associated with Edmondson grade), whereas the phosphoproteome clusters are not significantly associated with any other clustering (**Fig S5I**).

**Figure 5:**
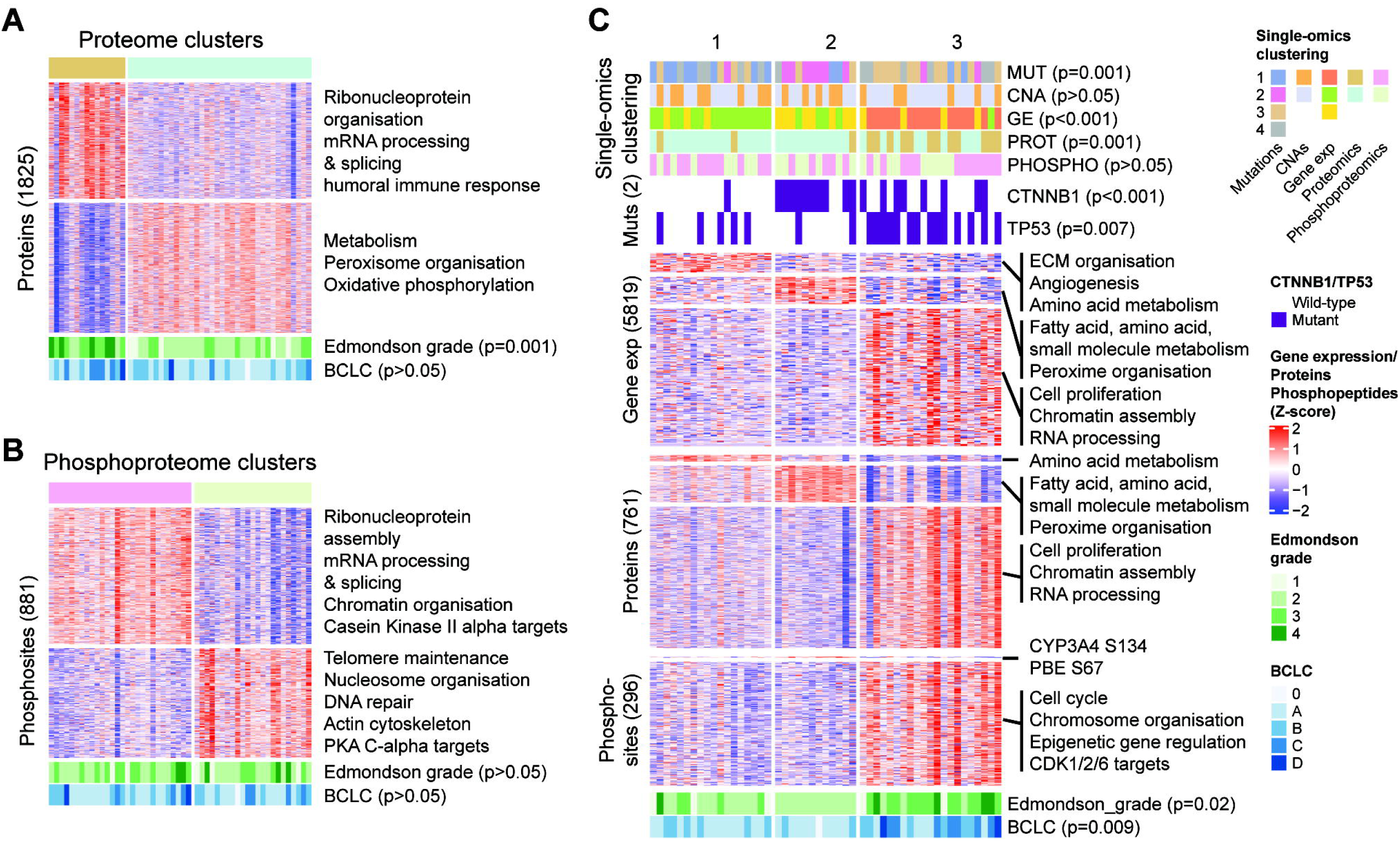
Integrated phosphoproteomic classification of HCC. **(A-B)** Unsupervised clustering of the **(A)** proteome and **(B)** phosphoproteome data using consensus non-negative matrix factorization. **(C)** Integrative clustering of the mutation, copy number alteration, transcriptome, proteome and phosphoproteome using the iCluster method. Copy number alterations not shown in the figure as no genomic region differed between clusters. See also Figures S5-6 and Tables S11-12.

We then asked whether an integrative clustering using all five layers of molecular information would provide further insight into the diversity of HCC. Using the iCluster method (Mo et al., 2018), we defined three robust clusters (Figure 5C), which are largely recapitulated using the algorithmically distinct similarity network fusion (SNF) approach (Wang et al., 2014) (**Figure S5J**). To investigate how these integrative clusters differ from each other, we compared them on the individual molecular levels (**Table S12**). In cluster 1, we observed an enrichment of genes involved in ECM organization, angiogenesis and amino acid metabolism on the mRNA level and amino acid metabolism on the protein level. In cluster 2, pathways related to fatty acid, amino acid and small molecule metabolism were enriched on both the mRNA and protein levels. In cluster 3, pathways related to cell proliferation, chromatin assembly and RNA processing were enriched on both the mRNA and protein levels, while on the phosphoprotein level, we observed pathways related to cell cycle, epigenetic gene regulation and activation of CDK1/2/6. Overall, the pathways enriched on the mRNA, protein and phosphoprotein levels are in accordance with the enrichment of *CTNNB1* mutations in cluster 2 and *TP53* mutations and higher Edmondson grade in cluster 3. No specific copy number alterations were found to be enriched in any of the subclasses. Compared to the single-omics clustering, the integrative clusters were associated with the mutation, transcriptome and proteome clusters (i.e. those that were associated with Edmondson grade) but not with the CNA or the phosphoproteome clusters (Figure 5C).

Finally, we asked whether the single-omics and integrative molecular subclasses are prognostic. Using univariate Cox proportional hazard analyses, transcriptome cluster 1 (increased cell proliferation and high Edmondson grade, p=0.037), proteome cluster 1 (ribonucleoprotein organization, mRNA processing and high Edmondson grade, p=0.022) and integrative cluster 3 (increased cell proliferation, epigenetic gene regulation, *TP53* mutation and high Edmondson grade, p=0.005) were associated with poor overall survival, while proteome cluster 2 (metabolism and low Edmondson grade, p=0.022) and integrative cluster 2 (metabolism, *CTNNB1* mutation and low Edmondson grade, p=0.015) were associated with improved overall survival (**Figure S6**). Given that BCLC is the primary prognostic indicator in HCC, we fit multivariate Cox proportional hazards models to evaluate whether the subclasses may be prognostic independent of BCLC. Here we observed that mutation cluster 2 (*CTNNB1*-mutant) was associated with good prognosis (p=0.025), while mutation cluster 3 (*TP53*-mutant) was associated with poor prognosis (p=0.034, **Figure S6**).

In summary, while molecular clustering of the proteome data largely recapitulated that of the transcriptome and histological grading, molecular clustering of the phosphoproteome data differed from that of other single-omics and integrative clustering.

## DISCUSSION

In this study, we performed a multiomic profiling of HCC biopsies across diverse etiologies and clinical stages. An integrated proteogenomic analysis revealed similar but also distinct biological processes, metabolic reprogramming and activation of signaling pathways on the different molecular levels. Our analysis provides novel insights into the molecular processes underlying HCCs.

Of the pathways altered in HCC, RNA processing was consistently upregulated and metabolic pathways were consistently downregulated on the transcriptome, proteome and phosphoproteome levels. Notably, metabolic pathways were enriched among hyperphosphorylated sites when we studied phosphorylation levels normalized by protein abundance. Conversely, genes related to translational regulation and oxidative phosphorylation were upregulated on the mRNA but not the protein level, whereas the complement and coagulation cascades were upregulated on the protein level but not the mRNA level. One could speculate that the increased transcription of genes related to translational regulation may be a compensatory mechanism for increased protein degradation. On the phosphoproteome level, altered protein phosphorylation was associated with pathways related to the cell cycle and the immune system, DNA repair and organization, as well as several oncogenic signaling pathways such as MAPK, PI3K/Akt/PTEN and FGFR. Indeed, KSEA revealed the increased activity of ERK1/2 and cell cycle-related kinases such as AURKA, CDK1/2/5/7, PLK1 and TTK.

Our proteogenomic analysis has also identified novel putative HCC driver genes and drug targets. In our analysis of genes and proteins that show positive correlation on the CNA-mRNA and mRNA-protein levels, we identified several candidate driver genes, such as *NUDCD1*, *SQLE*, *UBQLN4*, *ALYREF* and *CERS2*, involved in diverse processes including epithelial-to-mesenchymal transition (EMT), Wnt-β-catenin pathway regulation, transcriptional control, cholesterol biosynthesis and sphingolipid metabolism. In particular, *NUDCD1* (OVA66, 8q22.2 peak) and *SQLE* (8q24.13 peak) can both promote oncogenic transformation and/or cell proliferation and migration via major oncogenic signaling pathways such as PI3K/AKT, ERK1/2-MAPK and/or IGF-1R-MAPK (Rao et al., 2014a, 2014b; Sui et al., 2015). In the context of HCC, *SQLE* may be of particular interest given its role in cholesterol biosynthesis and that disrupted cholesterol homeostasis may be a hallmark in a subset of HCC (Jiang et al., 2019). Similarly, *CERS2*, a key component in sphingolipid metabolism, may also contribute to perturbations in sphingolipid synthesis that promote HCC development (Guri et al., 2017). Moreover, given that pathways related to RNA processing appear to be consistently altered on multiple molecular levels in HCC, genes involved in transcriptional control, such as *ALYREF* (Hautbergue et al., 2008; Hung et al., 2010; Stubbs and Conrad, 2015; Stubbs et al., 2012), are potential driver genes that warrant further study. Our analysis of the HCC phosphoproteome also revealed several targetable kinases with elevated activity in HCC, especially Aurora Kinase A and CDKs. Aurora Kinase A and CDK1/2 are classical cell cycle-related kinases whereas CDK5 regulates many biological processes, among which are angiogenesis and DNA damage response (Ehrlich et al., 2015; Liebl et al., 2010). Preclinical studies have shown that inhibitors of Aurora Kinase A or CDK1/2/5 are efficacious in HCC models (Benten et al., 2009; Dauch et al., 2016; Haider et al., 2013) and may act synergistically with sorafenib/regorafenib and chemotherapeutic agents (Ehrlich et al., 2015; Xu et al., 2019; Zhang et al., 2018).

The two most frequently mutated genes in HCC are *CTNNB1* and *TP53*. By identifying sites whose phosphorylation differed between mutant and wild-type HCCs, but were not associated with alterations in expression of the phosphorylated protein, we identified phosphorylation sites in proteins, such as ACF7, MAP1B, PDLIM2 and Par3-alpha, that may underpin the EMT phenotype frequently seen in *CTNNB1*-mutant HCCs. ACF7 (*MACF1*), part of the β-catenin destruction complex (Kimelman and Xu, 2006), is required for stabilizing Axin during its translocation from the cytoplasm to the cell membrane upon Wnt activation (Salinas, 2007). Wnt activation causes GSK3 kinase inactivation and inability to phosphorylate ACF7, and dephosphorylated ACF7 remains active and able to form necessary connections between microtubules and the actin cytoskeleton for migration to occur (Wu et al., 2011; Zaoui et al., 2010). Similarly, GSK3 kinase inactivation has been shown to decrease MAP1B phosphorylation facilitating microtubule assembly and migration processes. By contrast, PDLIM2, located at the actin cytoskeleton, is required for polarized cell migration. PDLIM2 phosphorylation leads to its stabilization in the cytoplasm, facilitating β-catenin activation and nuclear translocation (Cox et al., 2019). Par3-alpha (*PARD3*), a regulator of tight junction assembly at epithelial cell-cell contacts (Chen and Macara, 2005), changes the affinity for its interaction partners (Funahashi et al., 2013; Hurd et al., 2003; Khazaei and Püschel, 2009) upon phosphorylation, leading to loss of cell polarity and induction of migration. KSEA of *CTNNB1*-mutant HCCs also revealed increased activity of FGR, involved in immune response and cytoskeleton remodeling. Of note, a previous study identified ALDOA S36 phosphorylation as a molecular feature of *CTNNB1*-mutant HBV-associated HCCs (Gao et al., 2019), but we did not observe elevated ALDOA S36 phosphorylation in our cohort (FDR=0.98). On the other hand, KSEA of *TP53*-mutant HCCs identified increased activity of CDK1/2/5 that are key regulators of the cell cycle and mitotic spindle, which could in part explain the higher histological grade typically associated with *TP53* mutations. Our analysis also identified altered phosphorylation of proteins involved in lipid metabolism in *TP53*-mutant HCCs. p53 with a gain-of-function mutation has been reported to promote lipid synthesis via at least two mechanisms, by activating the SREBP transcription factors and the mevalonate pathway and by inhibiting AMPK to promote tumorigenesis (Freed-Pastor et al., 2012; Zhou et al., 2014).

Proteome and phosphoproteome classifications revealed two clusters each. Although for both classifications, one of the clusters was associated with overexpression or enhanced phosphorylation of proteins involved in ribonucleoprotein organization, as well as mRNA processing and splicing, there was little concordance between the two classifications. The lack of concordance of the phosphoproteome clusters was also seen with mutation and transcriptome clusters. As protein phosphorylation is highly dynamic and our profiling captures a snapshot of the tumor, phosphoproteomic data are inherently noisier than other types of molecular data. In fact, the lack of concordance of the phosphoproteome clusters was also seen when compared to integrative clustering. Integrative clustering, using the algorithmically distinct iCluster and SNF, identified three clusters that resembled the single-omics clusters by mutation status, transcriptome and proteome profiles but not those by copy number and phosphoproteome profiles. The three classes as defined by integrative clustering resemble the spectrum of Edmondson grade and BCLC. They also resemble the three proteome subclasses identified in two previous studies in HBV-related HCC, both of which described subclasses characterized by metabolic reprogramming, microenvironment dysregulation and cell proliferation (Gao et al., 2019; Jiang et al., 2019). In particular, the proliferation proteome subclasses in HBV-related HCC were found to be associated with tumor thrombus (Gao et al., 2019) and microscopic vascular invasion (Jiang et al., 2019). Here our integrative cluster 3 (increased cell proliferation, *TP53* mutation and high Edmondson grade) is associated with macro-vascular invasion (p=0.01, Fisher’s exact test) though the association is not statistically significant after multiple testing correction. As for the prognostic value of the multiomics clusters, integrative clusters 2 and 3 are associated with good and poor outcomes, respectively, although the associations were not significant after accounting for the difference in BCLC. By contrast, mutation clusters 2 (*CTNNB1*-mutant) and 3 (*TP53*-mutant) were associated with overall survival independent of BCLC. However, it should be noted that our cohort was accrued over a long period and clinical practice has changed significantly over the past decade, hence outcome data are inherently difficult to interpret.

In conclusion, our study provides a comprehensive analysis of the proteomic and phosphoproteomic landscape of HCCs, identifying proteome and phosphoproteome alterations underlying HCC.

## Acknowledgments

The project was funded by the ERC Synergy Grant 609883 (M.N.H. and M.H.H.). L.M.T., C.K.Y.N. and S.P. were supported by the Swiss Cancer League (KLS-3639-02-2015, KFS-4543-08-2018, KFS-4988-02-2020-R, respectively); L.M.T. was supported by AIRC grant number IG 2019 Id.23615, S.P. was supported by the Swiss National Science Foundation (PZ00P3_168165), the University of Basel (Research Fund Junior Researchers), Krebsliga Beider Basel (KLbB-4473-03-2018), the Theron Foundation, Vaduz (LI) and the Surgery Department of the University Hospital Basel. The funders had no role in study design, data collection, and analysis, decision to publish, or preparation of the manuscript. Data analysis was performed at sciCORE scientific computing center at the University of Basel.

## Author Contributions

L.M.T., M.N.H. and M.H.H. conceived and supervised the study; C.K.Y.N. performed the bioinformatic analysis; T.B. and M.H.H. performed the biopsy procedures; T.B., S.W. and M.H.H. provided the clinical information; C.E., M.S.M. and L.M.T. performed the histopathological assessment of tissue samples; X.W. and S.K. collected, processed and archived biopsy tissue; S.N., A.S., M.A.M. and S.W. processed the tissue samples and performed nucleic acid extraction for sequencing; E.D., M.C. and M.N.H. performed the proteome and phosphoproteome profiling and data processing; T.B. and A.S. helped with the phosphoproteome profiling and data processing; C.K.Y.N., E.D. and M.C.-L. wrote the manuscript, which was initially revised by S.P. and M.H.H. All authors have read and revised the manuscript.

## STAR METHODS

### HCC biopsy procedure and sample collection

Human tissues were obtained from patients undergoing diagnostic liver biopsy at the University Hospital Basel between 2008 and 2018. Written informed consent was obtained from all patients. The study was approved by the ethics committee of the northwestern part of Switzerland (Protocol Number EKNZ 2014-099). Ultrasound-guided needle biopsies were obtained from tumor lesion(s) and the liver parenchyma at a site distant from the tumor with a coaxial liver biopsy technique that allows taking several biopsy samples through a single biopsy needle tract as described (Nuciforo et al., 2018). Clinical disease staging was performed using the Barcelona Clinic Liver Cancer system (European Association for the Study of the Liver, 2018). Biopsies from multicentric tumors (i.e. genetically independent primary tumors), but not intra-hepatic metastases, were included. In total, 122 HCC biopsies and 115 non-tumoral tissues from 114 patients were included in the study **(Table S1)**, including 6 patients from whom 2 synchronous multicentric tumor biopsies and 1 patient from who, 3 multicentric tumor biopsies were obtained **(Figure S1)**. None of the patients had received systemic or locoregional therapies for liver cancer prior to biopsy. Two patients were treated with curative surgery or ablation and were biopsied after HCC recurrence was diagnosed by imaging.

As control, we used liver biopsies with normal histology obtained from 19 patients without HCC and with normal liver values **(Table S1)**. The biopsy procedure was as described above.

### Histopathological assessment

Diagnosis of HCC and histopathology evaluation were carried out on FFPE slides blindly by at least two board-certified hepatopathologists (CE, MSM and/or LMT) at the Institute of Pathology of the University Hospital Basel. Histopathologic grading was performed according to the Edmondson grading system (Edmondson and Steiner, 1954; Nuciforo et al., 2018). Hematoxylin & eosin (H&E) slides were reviewed to define the presence or absence of cirrhosis, underlying liver disease, cholestasis, vessel infiltration, necrotic areas, major growth pattern, cytological variants and special subtypes according to the guidelines by the World Health Organization (World Health Organization, 2010). Immunophenotypes were classified according to Chen et al (Chen and Mellman, 2017). Specifically, inflamed tumors are defined as tumors in which tumor infiltrating lymphocytes (TILs) are present in the tumor parenchyma in close proximity to tumor cells; immune-excluded are tumors in which TILs are present only in ≥10% of the tumor stroma and/or tumor margins is populated by lymphocytes located in the immediate vicinity of tumor cells; and immune-desert, in which less than 10% of the tumor stroma is populated by lymphocytes, and neither dense immune cell infiltrates nor immune cells are in contact with tumor cells.

### DNA and RNA extraction

Genomic DNA and total RNA from tumor and adjacent liver parenchyma were extracted using the ZR-Duet DNA and RNA MiniPrep Plus kit (Zymo Research) following the manufacturer’s instructions. Prior to extraction, biopsies were crushed in liquid nitrogen to facilitate lysis. Total RNA of 15 patients without HCC was extracted using Trizol (Thermo Fisher Scientific) according to the manufacturer’s instructions. Extracted DNA was quantified using the Qubit Fluorometer (Invitrogen). Extracted RNA was quantified using NanoDrop 2000 spectrophotometer (Thermo Fisher Scientific), and RNA quality/integrity was assessed with an Agilent 2100 BioAnalyzer using RNA 6000 Nano Kit (Agilent Technologies).

### Whole-exome sequencing and data processing

All 122 HCC biopsies and 115 non-tumoral tissues from 114 patients were subjected to whole-exome sequencing. Whole-exome capture was performed using the SureSelectXT Clinical Research Exome (Agilent Technologies) or SureSelect Human All Exon V6+COSMIC (Agilent Technologies) platforms according to the manufacturer’s guidelines. Sequencing was performed on an Illumina HiSeq 2500 at the Genomics Facility Basel according to the manufacturer’s guidelines. Paired-end 101-bp reads were generated. Tumor biopsies and non-tumoral biopsies were sequenced to median depths of 94.3 (range 16.4-140.0) and 49.4 (range 34.5-86.2), respectively (**Table S1**).

Sequence reads were aligned to the reference human genome GRCh37 using Burrows-Wheeler Aligner (BWA, v0.7.12/13) (Li and Durbin, 2009). Local realignment, duplicate removal and base quality adjustment were performed using the Genome Analysis Toolkit (GATK, v3.6) (McKenna et al., 2010) and Picard (http://broadinstitute.github.io/picard/, v2.4.1). Somatic single nucleotide variants (SNVs) and small insertions and deletions (indels) were detected using MuTect (v1.1.4) (Cibulskis et al., 2013) and Strelka (v1.0.15) (Saunders et al., 2012), respectively. We filtered out SNVs and indels outside of the target regions: those with variant allelic fraction (VAF) of <1% and/or those supported by <3 reads. We excluded variants for which the tumor VAF was <5 times that of the paired non-tumor VAF. We further excluded variants identified in at least two of a panel of 123 non-tumor samples, including the 115 non-tumor samples included in the current study, captured and sequenced using the same protocols using the artifact detection mode of MuTect2 implemented in GATK 3.6. Mutations affecting hotspot residues (Chang et al., 2016) were annotated. Significantly mutated genes were identified using MutsigCV (v1.4) (Lawrence et al., 2013) and OncodriveFML (Mularoni et al., 2016). Genes with q < 0.1 were considered significantly mutated. ‘Lollipop’ plots were generated using the ‘MutationMapper’ tool on the cBioPortal (Cerami et al., 2012). Mutual exclusivity and co-occurrence of significantly mutated genes were computed using one-sided Fisher’s exact test (p<0.05), where log_2_ odds ratio >0 indicates occurrence and log_2_ odds ratio <0 indicates mutual exclusivity. Tumor mutational burden was defined as the total number of somatic mutations (including synonymous and nonsynonymous point mutations and indels) in the coding region and splice sites.

Allele-specific CNAs were identified using FACETS (v0.5.5) (Shen and Seshan, 2016), which performs a joint segmentation of the total and allelic copy ratio and infers allele-specific copy number states. Somatic mutations associated with the loss of the wild-type allele (i.e., loss of heterozygosity [LOH]) were identified as those for which the lesser (minor) copy number state at the locus was 0. All mutations on chromosome X in male patients were considered to be associated with LOH. Copy number states were collapsed to the gene level based on the median values to coding gene resolution based on all coding genes retrieved from the Ensembl (release GRCh37.p13). Genes with total copy number greater than gene-level median ploidy were considered gains; greater than ploidy + 4, amplifications; less than ploidy, losses; and total copy number of 0, homozygous deletions. Fraction of genome altered was computed as the fraction of genes with amplification, gain, loss or deletion. Tumors with >5% of the genome at copy number 0 (homozygous deletions, 5 tumors) were excluded from the identification of homozygous deletions and from the computation of fraction of genome altered. Significant focal copy number alterations were identified from segmented data for all 122 tumor biopsies using GISTIC2.0 (v2.0.23) (Mermel et al., 2011).

### RNA-sequencing and data processing

RNA-seq library prep was performed with 200 ng total RNA using the TruSeq Stranded Total RNA Library Prep Kit with Ribo-Zero Gold (Illumina) according to manufacturer’s specifications. Single-end 126-bp sequencing was performed on an Illumina HiSeq 2500 using v4 SBS chemistry at the Genomics Facility Basel according to the manufacturer’s guidelines. Primary data analysis was performed with the Illumina RTA version 1.18.66.3.

Sequence reads were aligned simultaneously to the human reference genome GRCh37, HBV strain ayw genome (NC 003977.2), and HCV genotype 1 genome (NC 004102.1) by STAR (v2.5.2a) (Dobin et al., 2013) using the two-pass approach. Median numbers of reads aligning to the human genome were 52.2 million (range 37.4 - 115.1 million) and 63.5 million (range 52.5 -82.2 million) for the HCC and normal liver biopsies, respectively (**Table S1**).

Transcript quantification was performed using RSEM (v1.2.31) (Li and Dewey, 2011). Gene-level expected counts were upper-quartile-normalized to 1000. For downstream analysis, we computed the log_2_-fold-changes of normalized RSEM gene counts between tumors and the median of 15 normal livers.

### Biopsy sample preparation for proteomics, protein extraction and digestion

Fresh liver biopsies from 51 HCC and 11 normal livers were immediately snap-frozen in liquid nitrogen and processed as previously described (Dazert et al., 2016). The average time from removing the biopsy from the liver to freezing took about 2 min. In brief, for protein extraction, each frozen biopsy was crushed in an in-house constructed metal mortar cooled on dry ice into a fine powder (cryogenic grinding) and transferred into a cooled 1.5 ml tube containing 150 - 400 ml lysis buffer (50 mM Tris-HCl pH 8.0, 8M urea, 150 mM NaCl, 1 mM PMSF, Complete Mini Protease Inhibitors (Sigma-Aldrich), PhosSTOP Phosphatase Inhibitors (Sigma-Aldrich)). The biopsy lysate was vigorously vortexed for 5 min, rotated for 1h at 4°C and sonicated twice in a VWR Ultrasonic cleaner bath (USC300T) for 1 min. Next, the lysate was centrifuged for 10 min at 15°C at 14.000 rpm and supernatant was removed and stored. Next, 50 μl of fresh lysis buffer were added and the sample was homogenized with a Teflon pestle in a hand homogenizer (Pellet Pestle Motor, Kontes/Kimble, USA) at maximum speed on ice twice for 1 min. Samples were centrifuged for 10 min at 15°C at 14.000 rpm and supernatant was removed and pooled with previous one. Protein concentration was measured with a Bradford assay. Next, proteins were reduced with 10 mM DTT for 1h at 37 °C and alkylated with 50 mM iodoacetamide for 30 min at RT in the dark, both with gentle shaking. Urea concentration was lowered to 4 M with 50 mM Tris-HCl, pH 8.0. Lysates were digested with two rounds of endoproteinase LysC (Wako) at a 1:100 enzyme-to-protein ratio at 37°C for two hours. Next, the urea concentration was lowered to 1 M. Lysates were digested with two rounds of trypsin (Sigma): 1:50 ratio overnight and 1:100 ratio for 2 h at 37°C. Digestion was stopped with TFA to a final concentration of 0.5%. Digests were centrifuged for 2 min at 1,500 g and desalted on a C18 SepPak cartridge (50mg column for up to 2.5mg peptide load capacity) (Waters) or C18 Macrospin/Microspin cartridge (Waters). Peptide concentration was estimated at 280nm, aliquots were taken and peptides were dried in the SpeedVac.

### Peptide fractionation for proteome of human HCC biopsies

Human HCC biopsies were measured by sequential window acquisition of all theoretical mass spectra (SWATH), in which data-independent acquisition is coupled with spectral library match (Gillet et al., 2012). From each biopsy, 30 μg of peptides were used for SWATH analysis and 100 μg of peptides were used for library preparation. The biopsies from the 11 patients with healthy livers were measured individually and also as a pool. This pool was measured as a reference several times over the course of the project to account for potential batch effects. Ten biopsy samples were measured together as one batch of samples on the same capillary column. For library generation, 100 μg of peptides from each of the 10 biopsies of one batch were pooled together and subjected to high-pH fractionation with a total of 1mg of peptide injected by 5 individual injections of 200 μg. Peptides were fractionated by high-pH reversed phase separation using a XBridge Peptide BEH C18 column (3,5 μm, 130 Å, 1mm x 150 mm, Waters) on an Agilent 1260 Infinity HPLC system. Peptides were loaded on column in buffer A (ammonium formate (20 mM, pH 10) in H2O) and eluted using a two-step linear 60 min gradient from 2% to 50% (v/v) buffer B (90% acetonitrile / 10% ammonium formate (20 mM, pH 10) at a flow rate of 42 μl/min. Elution of peptides was monitored with a UV detector (215 nm, 254 nm). A total of 36 fractions were collected and subsequently pooled into 12 fractions using a post-concatenation strategy as previously described (Wang et al., 2011). Peptides were dried in a SpeedVac, resuspended in 0.1% formic acid (mobile phase A) and OD was measured. Twenty μg of each fraction were used for library measurements.

### SWATH analysis and library preparation

The biopsy samples were analyzed on a Thermo Fisher QExactive Plus instrument coupled to an Easy nLC 1000. For SWATH analysis of the biopsy samples, 1.1 μg was injected on column including 10% of iRT peptide mix (HRM kit Ki-3003, Biognosys, Zurich, Switzerland). For library generation, 2 μg of each high pH fraction including 10% of iRT peptide mix (HRM kit Ki-3003, Biognosys, Zurich, Switzerland) were injected on column. Proteomes were analyzed by capillary LC-MS/MS using a homemade separating column (0.075 mm × 38 cm) packed with Reprosil C18 reverse-phase material (2.4-μm particle size; Dr. Maisch). The solvents used for peptide separation were 0.1% formic acid (solvent A) and 0.1% formic acid and 80% AcCN in water (solvent B). Two microliters of sample were injected. A linear gradient from 0–40% solvent B in solvent A in 190 min was delivered with the nano pump at a flow rate of 200 nL/min. After 190 min, solvent B was increased to 95% in 5 min. The eluting peptides were ionized at 2.5 kV. Singly charged ions and ions with unassigned charge state were excluded from triggering MS2 events. For SWATH measurements, one Full MS-SIM scan at resolution of 70.000 was followed by 40 mass windows of dynamic size ranging from 400 to 1600 m/z with 4 kDa overlap at a resolution of 17.500. For library measurements, the mass spectrometer was operated in data-dependent mode and the precursor scan was done in the Orbitrap at 70,000 resolution. A top-20 method was run. For SWATH analysis and library generation, samples were injected in triplicates.

### SWATH data analysis

The library was generated with MaxQuant (version 1.5.1.2) (Cox and Mann, 2008) using the default settings except that the mass tolerance of the instrument was set to 10 ppm and the minimal ratio count for quantification was set to 1. The Uniprot SwissProt database (17^th^ August 2015) including the iRT fusion peptide was used for the searches. All library measurements were pooled into one MaxQuant analysis to generate one general HCC library. The raw SWATH MS runs of the individual biopsies were converted using the HTRMS converter (Biognosys). The converted SWATH runs were analyzed with Spectronaut X (Version 12.0.20491.20.29183) (Biognosys) using the default settings and searched against our in-house generated general HCC library.

### Proteome analysis

Raw protein-group based data were exported from Spectronaut and imported into FileMakerPro Advanced (Version FMP18) for further data processing. The raw intensities of the triplicates were averaged and the mean values transformed by the logarithm to the base 2. Next, the values were normalized by median subtraction. To account for potential batch effects, the log_2_ median-subtracted intensities of each biopsy were normalized to the mean intensity of all measured runs of the pool of healthy liver tissue. The proteome of the patient biopsies was continuously measured over a time frame of 2 years. Throughout this time period also aliquots of the pooled healthy sample were measured. All measured runs of the pooled healthy sample were therefore averaged for normalization.

We obtained data for 6167 proteins that were quantified (always against healthy liver) in at least one run in at least one HCC (**Table S1**), 5612 proteins quantified in at least 26 HCCs and 1997 in at all 51 HCCs. Starting with the 6167 proteins quantified in at least one HCC, we removed proteins for which data were missing from >50% of the samples (50% to enable sufficient data for imputation), resulting in 5631 proteins for further analysis. Data imputation using nearest neighbor averaging was performed using the ‘impute.knn’ function from the ‘impute’ R package (v1.64.0).

### Phospho-proteome analysis

Peptide samples were enriched for phosphorylated peptides using Fe(III)-IMAC cartridges on an AssayMAP Bravo platform as recently described (Post et al., 2017). We used an input peptide amount of approx. 165 μg for the phosphoenrichment. For 3 biopsies, input to phosphoenrichment was slightly reduced but was accounted for by Progenesis/SafeQuant (see below). These 3 samples did not show an outlier pattern in terms of the quantified phosphorylation sites after data processing and were included in subsequent analyses. The μRPLC-MS system was set up as described previously (Ahrné et al., 2016). Chromatographic separation of peptides was carried out using an EASY nano-LC 1000 system (Thermo Fisher Scientific), equipped with a heated RP-HPLC column (75 μm x 37 cm) packed in-house with 1.9 μm C18 resin (Reprosil-AQ Pur, Dr. Maisch). Dried phosphopeptides were resuspended in 20 μl of 0.1% formic acid and 3 μl of the sample were injected per triplicate LC-MS/MS run. Samples were analyzed using a linear gradient ranging from 95% solvent A (0.15% formic acid, 2% acetonitrile) and 5% solvent B (98% acetonitrile, 2% water, 0.15% formic acid) to 30% solvent B over 90 minutes at a flow rate of 200 nl/min. Mass spectrometry analysis was performed on a Q-Exactive HF mass spectrometer equipped with a nanoelectrospray ion source (both Thermo Fisher Scientific). Each MS1 scan was followed by high-collision-dissociation (HCD) of the 10 most abundant precursor ions with dynamic exclusion for 20 seconds. Total cycle time was approximately 1 s. For MS1, 3e6 ions were accumulated in the Orbitrap cell over a maximum time of 100 ms and scanned at a resolution of 120,000 FWHM (at 200 m/z). MS2 scans were acquired at a target setting of 1e5 ions, accumulation time of 50 ms and a resolution of 30,000 FWHM (at 200 m/z). Singly charged ions and ions with unassigned charge state were excluded from triggering MS2 events. The normalized collision energy was set to 27%, the mass isolation window was set to 1.4 m/z and one microscan was acquired for each spectrum. The samples were measured in triplicates. The acquired raw-files were imported into the Progenesis QI software (v2.0, Nonlinear Dynamics Limited), which was used to extract peptide precursor ion intensities across all samples applying the default parameters. The generated mgf-files were searched using MASCOT against a decoy database containing normal and reverse sequences of the predicted SwissProt entries of Homo sapiens (www.ebi.ac.uk) and commonly observed contaminants generated using the SequenceReverser tool from the MaxQuant software (version 1.0.13.13). The search criteria were set as follows: full tryptic specificity was required; 3 missed cleavages were allowed; carbamidomethylation (C) was set as fixed modification; oxidation (M) and phosphorylation (STY) were applied as variable modifications; mass tolerance of 10 ppm (precursor) and 0.02 Da (fragments). The database search results were filtered using the ion score to set the false discovery rate (FDR) to 1% on the peptide and protein level, respectively, based on the number of reverse protein sequence hits in the datasets. The relative quantitative data obtained were normalized and statistically analyzed using our in-house script SafeQuant (Ahrné et al., 2016). Here the gMin algorithm was chosen. Afterwards, data were imported into FileMakerPro Advanced (Version FMP18) for further data processing. Imputed values were excluded and data were median subtracted per biopsy.

### Processing of phospho-proteome data

Our SafeQuant in house script generated phospho-peptide centric quantifications. In order to generate quantitative data for single phosphorylation sites, peptides with more than one phosphorylation site were deconvoluted. As a next step all intensities assigned to a single phosphorylation site were added up to generate one cumulative intensity per phosphorylation site. The raw intensities of the triplicates were averaged and the mean values transformed by the logarithm to the base 2. Next, the values were normalized by median subtraction. The phospho-enrichments were performed and measured in three batches due to the limitation of the number of MS runs that can be performed using the same capillary column. In each batch also an aliquot of the pooled healthy sample was enriched and measured. Normalization to the pooled healthy sample was then performed batch-wise to the pooled healthy sample enriched and measured at the same time with the same batch. Localization probabilities of each phosphorylation site were determined per batch using ScaffoldPTM (Version 3.2.0) (Proteome Software) and the maximum observed localization probability was assigned to each phosphorylation site. Only phosphorylation sites with a minimum localization probability of 50% were taken into account.

We obtained data for 12205 phosphorylation sites (in 4230 proteins) that were quantified (always against healthy liver) in at least one HCC (**Table S1**), 9606 (3816) quantified in at least one HCC with >99% localization probability, 7911 (3160) quantified in at least 26 HCCs, 6403 (2856) quantified in at least 26 HCCs with >99% localization probability, 4112 (2031) quantified in all 51 HCCs and 3439 (1837) quantified in at all 51 HCCs with >99% localization probability. Starting from the 12205 phosphorylation sites, we removed proteins for which data were missing from >50% of samples (50% to enable sufficient data for imputation), resulting in 7893 phosphorylation sites from 3156 proteins for further analysis. Since data were generated over three batches, we corrected for the batch effect using the ‘removeBatchEffect’ function in the *edgeR* R package (Robinson et al., 2010). Data imputation using nearest neighbor averaging was performed using the ‘impute.knn’ function from the *impute* R package. To normalize for overall protein levels, we computed the difference between the log_2_-fold-changes of phosphorylation site levels between tumors and the normal livers and the log_2_-fold-changes of protein levels between tumors and the normal livers, for proteins detected by both technologies.

### Differential expression analysis

For transcriptome data, differential expression analysis using the ‘edgeR’ package (v3.32.0) (Robinson et al., 2010) between samples from a given class vs all other samples using raw RSEM expected counts as input. Specifically, normalization was performed using the “TMM” (weighted trimmed mean) method (Robinson and Oshlack, 2010) and differential expression was assessed using the quasi-likelihood F-test, adjusted for multiple testing using Benjamini and Hochberg’s method. For proteome and phosphoproteome data, differential expression analysis was performed using the ‘limma’ package (v3.46.0) (Ritchie et al., 2015), using the log_2_-fold-changes of protein/phosphorylation site levels between tumors and the normal livers. *limma* fits a linear model to compute the moderated t-statistics using a Bayesian model and adjusts the p-values for multiple testing using Benjamini and Hochberg’s method. Genes, proteins and phosphorylation sites with adjusted p≤0.05 were considered differentially expressed.

### Pathway analysis

Pathway analysis (over-representation analysis and gene set enrichment analysis (GSEA)) was performed using the ‘clusterProfiler’ (v3.18.0) and ‘ReactomePA’ (v1.34.0) packages (Yu and He, 2016; Yu et al., 2012) for KEGG/Reactome pathways and Gene Ontology biological processes subset. For proteome and phosphoproteome data, the corresponding sets of detected proteins were used as background for overrepresentation tests. p≤0.05 was considered statistically significant. Pathway analysis results were represented as barplots, dotplots or enrichment maps.

### Kinase-substrate enrichment analysis (KSEA)

KSEA (Casado et al., 2013) was performed using the ‘KSEAapp’ R package (v0.99.0) (Wiredja et al., 2017) using NetworKIN.cutoff=5, using the log fold-change and p-values computed from differential expression analysis (see **Differential expression analysis**) of unimputed phosphorylation site levels and unimputed phosphorylation site levels normalized by overall protein levels (see **Processing of phospho-proteome data**) as input.

### Analysis of dysregulated genes/proteins and pathways in HCC

For the assessment of dysregulated genes and proteins in HCC, we performed differential expression analysis between HCCs and normal livers (see **Differential expression analysis**) for transcriptome and proteome data. To compare the dysregulated genes/proteins between transcriptome data and proteome data, Uniprot accessions were mapped to Ensembl gene IDs, resulting in 5490 comparable genes/proteins. Dysregulated pathways were identified using a quadrant analysis, by performing over-representation tests (see **Pathway analysis**).

### CNA-mRNA-protein correlation

Correlation was performed using segmented log ratio for CNA, and the log_2_-fold-changes of protein levels between tumors and the median of normal livers for mRNA and protein data. Uniprot accessions were mapped to Ensembl gene IDs. For the CNA-mRNA correlation, 15272 genes were included. For the mRNA-protein correlation, 5481 genes were included. CNA-mRNA and mRNA-protein correlations were assessed using Spearman correlation tests. To assess the enrichment of genes within significant focal copy number alterations defined by GISTIC, genes were ranked according to Spearman correlation coefficient for GSEA analysis using the *clusterProfiler* package (Yu et al., 2012). p-value ≤ 0.05 was considered statistically significant.

### Analysis of dysregulated phosphorylation sites in HCC

For the assessment of dysregulated phosphorylation sites in HCC, we performed differential expression analysis between HCCs and normal livers (see **Differential expression analysis**) for phosphorylation site levels with and without normalization by protein level. Significantly regulated phosphorylation sites (adjusted p < 0.05 and |logFC|≥2) were used for pathway analysis using over-representation tests (see **Pathway analysis**), separately for up- and down-regulated phosphorylation sites, as well as for up- and down-regulated phosphorylation sites together. KSEA was also performed to identify differential kinase activity by computing the differential expression between HCCs and normal livers (see **Kinase-substrate enrichment analysis**).

### Phosphoproteomic analysis for *CTNNB1* and *TP53* mutations

Transcriptomic, proteomic and phosphoproteomic signatures of *CTNNB1* and *TP53* mutations were identified by differential expression analysis of the HCCs (see **Differential expression analysis**), by fitting a single model incorporating the mutation status of both genes. To identify phosphorylation sites associated with mutations in these two genes but were not associated with differences on the protein levels, we identified phosphorylation sites whose p-values were within the most extreme 10th percentile while the p-values of the corresponding proteins were within inter-quartile range. These phosphorylation sites were then subjected to pathway analysis by over-representation tests (see **Pathway analysis**). KSEA was also performed to identify differential kinase activity by computing the differential expression between mutant and wild-type HCCs (see **Kinase-substrate enrichment analysis**).

### Single-omics clustering

Identification of tumor subclasses based on somatic non-synonymous mutations was performed using oncosign (v1.0) (Ciriello et al., 2013) and Network-Based Stratification (pyNBS, downloaded on 4th June 2020) (Hofree et al., 2013; Huang et al., 2018). Significantly mutated genes identified using MutsigCV (Lawrence et al., 2013), as well as genes identified as significantly mutated in HCC in at least 2 of Martincorena et al(Martincorena et al., 2018), Schultz et al (Schulze et al., 2015), Fujimoto et al (Fujimoto et al., 2016), Bailey et al (Bailey et al., 2018) (excluding *TERT*) and mutated (non-synonymous) in at least 3 tumor samples were included for the clustering.

Identification of tumor subclasses based on copy number alterations was performed using consensus k-means clustering and consensus hierarchical clustering using the ‘ConsensusClusterPlus’ R package (v1.54.0) (Wilkerson and Hayes, 2010), using gene-level amplification, gain, neutral, loss and deletion status as input. 117 tumors were included, excluding the 5 for which copy number gain/loss status could not be determined (see **Whole-exome sequencing and data processing**). For both clustering methods, Euclidean distance was used as the distance metric, and up to 10 clusters were evaluated over 100 subsamples. Hierarchical clustering was performed using the “ward.D2” method.

Identification of tumor subclasses based on transcriptome, proteome and phosphoproteome subclasses was performed using consensus nonnegative matrix factorization (NMF) and consensus k-means clustering using the ‘CancerSubtypes’ (v1.16.0) and the ‘ConsensusClusterPlus’ (v1.54.0) R packages (Wilkerson and Hayes, 2010; Xu et al., 2017), respectively. Log2-fold-change between tumors and the median of normal livers were used as input. For transcriptome and phosphoproteome clustering, features with standard deviation ≥2 across the tumors were included for clustering, resulting in 1370 and 1024 features, respectively. For proteome clustering, features with standard deviation ≥1 across the tumors were included, resulting in 1083 features. For consensus NMF, up to 10 clusters were evaluated over 50 NMF runs. For consensus k-means clustering, 1-Spearman correlation coefficient was used as the distance metric, and up to 10 clusters were evaluated over 100 subsamples.

Robustness of the subclasses was assessed by downsampling to 70%, 80% or 90% of samples over 20 runs, reclustering the reduced dataset, and calculating the adjusted Rand index compared to the full dataset. Cluster quality was assessed by Silhouette widths (except for mutation subclasses). The final number of clusters was chosen on the basis of the Silhouette widths and adjusted Rand index of the full dataset and for the 20 iterations of the downsampled datasets.

For mutation subclasses, the enrichment of mutated genes was assessed using a chi-squared test across all clusters and using Fisher’s exact tests comparing a given cluster to all other clusters. For CNA subclasses, the enrichment of copy number-altered genes was assessed using Mann-Whitney U tests, adjusted for multiple testing using Benjamini and Hochberg’s method. Genes with adjusted p≤0.05 were considered statistically significant. For transcriptome, proteome and phosphoproteome subclasses, over-expressed features were identified by differential expression analysis between all samples in a given class and all other samples (see **Differential expression analysis**) followed by pathway analysis (see **Pathway analysis**). KSEA was also performed for the phosphoproteomics subclasses (see **Kinase-substrate enrichment analysis**). Figures were generated using the ‘ComplexHeatmap’ R package (v2.6.2) (Gu et al., 2016).

### Integrative clustering

Integrative clustering was performed for the 51 HCCs for which data were available for all data types using the ‘iClusterBayes’ function, which fits a Bayesian latent variable model that generates an integrated cluster assignment based on joint inference across data types, implemented in the ‘iClusterPlus’ R package (v1.26.0) (Mo et al., 2018) and the similarity network fusion (SNF) method (Wang et al., 2014), which constructs a fusion patient similarity network by integrating the patient similarity obtained from each of the genomic data types, as implemented in the ‘SNFtool’ (v2.3.0) and the ‘CancerSubtypes’ R packages (Wang et al., 2014; Xu et al., 2017). As input data, significantly mutated genes identified using MutsigCV (Lawrence et al., 2013), as well as genes identified as significantly mutated in HCC in at least 2 of Martincorena et al (Bailey et al., 2018; Martincorena et al., 2018), Schultz et al (Schulze et al., 2015), Fujimoto et al (Fujimoto et al., 2016), Bailey et al (Bailey et al., 2018) (excluding *TERT*) and mutated (non-synonymous) in at least 2 samples were included as mutational data. CNA data were included as collapsed copy number regions, constructed using the ‘CNregions’ function in the ‘iClusterPlus’ R package to reduce the segmented logR ratio, as recommended in the package vignette, resulting in 927 features for clustering. For transcriptome and phosphoproteome data, features with standard deviation ≥2 across the tumors were included for clustering, resulting in 1646 and 1024 features, respectively. For proteome data, features with standard deviation ≥1 across the tumors were included, resulting in 1083 features. Transcriptome, proteome and phosphoproteome data were z-score-transformed prior to clustering. For both clustering, up to 10 clusters were evaluated.

Robustness of the subclasses was assessed by downsampling to 70%, 80% or 90% of samples over 20 runs, reclustering the reduced dataset, and calculating the adjusted Rand index compared to the full dataset. For iClusterBayes, the final number of clusters was chosen on the basis of the Bayesian Information Criterion, the deviance ratio (interpreted as percent explained variation) and adjusted Rand index between the full dataset and the 20 iterations of the downsampled datasets. For SNF, cluster quality was assessed by Silhouette widths and the final number of clusters was chosen on the basis of the Silhouette widths and adjusted Rand index of the full dataset and for the 20 iterations of the downsampled datasets.

The identification of enriched features were performed as per single-omics clustering for the corresponding data type. Figures were generated using the ‘ComplexHeatmap’ R package (v2.6.2) (Gu et al., 2016).

### Statistical Analysis

Principal component analysis (PCA) of transcriptome, proteome and phosphoproteome data was performed using ‘prcomp’ from the *stats* R library. For transcriptome data, the upper-quartile-normalized RSEM values were used as input. For proteome and phosphoproteome data, the log_2_-fold-changes of protein/phosphorylation site abundance between tumors and the median of normal livers were used. Intra-group variability from PCA was computed as the pairwise Euclidean distance between samples of the same Edmondson grade using all principal components. Distance to normal livers was computed as the Euclidean distance between a given HCC sample to the median of normal livers using all principal components.

Statistical analyses of the clinicopathological variables were performed in R version 4.0.3. Comparisons of ordinal variables (e.g. BCLC, Edmondson grade, number of tumors) were performed using Mann-Whitney U tests. Comparisons of categorical variables (e.g. immunophenotype, presence of metastasis) were performed using Fisher’s exact tests or chi-squared tests. Comparisons of numerical variables (e.g. tumor mutational burden) were performed using Mann-Whitney U or Kruskal-Wallis tests. Correction for multiple testing was performed using the Benjamini-Hochberg method.

The association of overall survival and molecular subclasses was performed using Cox proportional-hazards model, including BCLC stage as a covariate. Overall survival was defined as the time interval between the diagnosis of HCC to death. Individuals who were lost-to-followup or had undergone liver transplantation were considered censored. For patients with >1 biopsy included in the study, only one biopsy was considered if all biopsies were of the same molecular subclass and patients excluded from overall survival analysis if the biopsies were of multiple molecular subclasses. All statistical tests were two-sided unless otherwise indicated, and p≤0.05 was considered statistically significant.

### Resource Availability

Lead Contact: Further information and requests for resources should be directed to and will be fulfilled by the Lead Contact, Markus H Heim.

Data Availability: The sequencing datasets generated during this study are available at European Genome-phenome Archive under accessions EGAS00001005073 (whole-exome sequencing) and EGAS00001005074 (RNA-sequencing).

## SUPPLEMENTAL INFORMATION TITLES AND LEGENDS

**Figure S1:** Heatmaps illustrate the variant allele fractions (shades of blue according to the color key, gray indicates absence) of the somatic mutations identified in the 7 patients for whosm >1 tumor biopsy was included in the study. Non-synonymous mutations in HCC driver genes are labeled. To the right of the heatmaps are genome-wide copy number plots of the tumor biopsies. In the copy number plots, segmented Log2 ratios (y-axis) were plotted according to their genomic positions (x-axis). Alternating blue and gray demarcate the chromosomes. Related to Table 1.

**Figure S2: (A)** Oncoprint showing the somatic mutational landscape of HCC biopsies, stratified by the availability of proteome and phosphoproteome profiling. Significantly mutated genes in the current cohort (*CTNNB1*, *CDKN1A*, *TP53*, *ALB*, *ARID1A*, *GPAM, AXIN1)* and six additional HCC driver genes (previously reported in at least 2 studies and mutated in at least 3 biopsies in this study) are included. Barplot above the oncoprint shows the total number of somatic mutations in each biopsy. Percentages to the left of the oncoprint show the fraction of biopsies harboring somatic mutations in a given gene. Barplot to the right of the oncoprint shows the total number and type of mutations identified in a given gene. The type of mutations is color-coded according to the legend. (**B**) Oncoprint showing the copy number profiles of HCC biopsies, stratified by the availability of proteome and phosphoproteome profiling. Significantly altered regions as defined by GISTIC2 on the entire cohort are shown. Copy number status was defined by GISTIC2. (**C**) Principal component analysis plot of gene expression of HCC biopsies (filled circles) and normal liver biopsies (open circles), colored by the availability of proteome and phosphoproteome profiling. Related to Table 1.

**Figure S3: (A)** Volcano plot of the −log10(adjusted p-value) against the log fold-change (logFC) of the differentially regulated phosphorylation sites normalized by overall protein levels (’normalized phosphorylation sites’) in HCC compared to normal livers. Dots are colored by logFC. Vertical dotted lines indicate |logFC|=2 and horizontal dotted lines indicate adjusted p-value=0.05. **(B)** Dot plot illustrating selected enriched Reactome pathways according to gene set enrichment analysis (GSEA) from the differential expression analysis in (**A**). NES: normalized enrichment score. **(C)** Top barplot showing the enrichment z-score of the kinases with significantly up- or downregulated kinase activity in a kinase-substrate enrichment analysis (KSEA) comparing normalized phosphorylation sites in HCC to normal livers. In the bubble plot below, the phosphorylation site substrates are shown in rows, where red and blue dots indicate that the phosphorylation site is up- and downregulated, respectively. The size of the dots is proportional to the log_2_ fold-change of the phosphorylation site. Phosphorylation sites with at least a 5-fold difference between HCCs and normal livers are shown. For kinases with <3 substrates with at least a 5-fold difference, the top three substrates with the highest |logFC| are shown. Related to Figure 3.

**Figure S4: (A)** Oncoprint showing the somatic mutational landscape of HCC. Significantly mutated genes in the current cohort (*CTNNB1*, *CDKN1A*, *TP53*, *ALB*, *ARID1A*, *GPAM, AXIN1*, in bold) and six additional HCC driver genes (previously reported in at least 2 studies and mutated in at least 3 biopsies in this study) are included. Barplot above the oncoprint shows the total number of somatic mutations in each biopsy. Percentages to the left of the oncoprint show the fraction of biopsies harboring somatic mutations in a given gene. Barplot to the right of the oncoprint shows the total number and type of mutations identified in a given gene. The type of mutations is color-coded according to the legend. **(B)** Lollipop plot showing the distribution of the *GPAM* mutations along the protein, with the mutations colored according to the color key in **(A)**. **(C)** Bubble plot showing association between mutation status and clinicopathological parameters. Size of the circles is proportional to −log10(p-value) and blue circles indicate statistically significant associations. Statistical analyses were performed by Fisher’s exact or Chi-squared tests. (**D**) Forest plot showing multivariate Cox proportional-hazards model of overall survival according to the mutation status of HCC driver genes and BCLC clinical staging. Related to Figure 4.

**Figure S5:** For the clustering of HCC biopsies based on somatic mutations, we used OncoSign (primary) and pyNBS (alternative). For CNA, we used consensus k-means clustering (primary) and consensus hierarchical clustering (alternative). For the remaining data types, we used consensus nonnegative matrix factorization (NMF, primary) and consensus k-means clustering (alternative). We assessed the mean Silhouette width (cluster quality) and adjusted Rand index (concordance with the classes derived from the full dataset) by subsampling 70%, 80% and 90% of the samples. (**A**, left) Boxplot showing adjusted Rand index between the classes derived from OncoSign (single-omics clustering using mutation data) using the full data set and the classes derived from downsampled data. Downsampling (70%, 80% and 90% of the samples) was performed over 20 iterations. (right) Concordance between clusters derived from OncoSign and pyNBS. (**B-E**, from left) Mean silhouette widths against k (the number of clusters), mean silhouette widths against k from clustering using downsampled datasets (70%, 80% and 90% of the samples, 20 iterations of downsampling), adjusted Rand index between the classes derived from the full dataset and the classes derived from subsampled data using the primary clustering method, and concordance between the primary and alternative clustering methods. Statistical comparisons were performed by Fisher’s exact or chi-squared tests. **(F)** Clusters derived from OncoSign based on the significantly mutated genes found in this study and in previous studies (see methods). **(G)** Clusters derived from copy number alterations by consensus K-means clustering. **(H)** Clusters derived from gene expression by consensus NMF. **(I)** Concordance between single-omics clusters. Statistical comparisons were performed by Fisher’s exact or chi-squared tests. **(J**, top) Assessment of the Bayesian Information Criterion and the deviance ratio (interpreted as percent explained variation) from integrative clustering using iClusterBayes. (bottom left) Boxplot showing adjusted Rank index between the classes derived from integrative clustering using the full data set and the classes derived from downsampled data. Downsampling (70%, 80% and 90% of the samples) was performed over 20 iterations. (bottom right) Concordance between clusters derived from iClusterBayes and SNF. Statistical comparison was performed using a chi-squared test. Related to Figure 5.

**Figure S6:** Forest plots from Cox proportional hazards analyses. (Left) Univariate analysis for each single-omics and integrative clusters. (Right) Multivariate analysis for each single-omics and integrative clusters incorporating BCLC clinical stage as a covariate. Related to Figure 5.

**Table S1:** Clinicopathological parameters of the cohort, as well as technical details of molecular profiling. Related to Table 1.

**Table S2:** List of enriched Reacome, KEGG and Gene Ontology biological processes for deregulated pathways on the mRNA and protein levels. Related to Figure 1F.

**Table S3:** List of enriched Reactome/KEGG pathways/GISTIC2 peaks among genes with high CNA-mRNA expression correlation and genes with high mRNA-protein expression correlation. Related to Figure 2C-D.

**Table S4:** List of genes with high (rho≥0.5) CNA-mRNA and mRNA-protein correlations. Genes within GISTIC2 peaks and GISTIC2 peaks enriched among genes with high CNA-mRNA correlation are annotated. Related to Figure 2D.

**Table S5:** List of dysregulated phosphorylation sites in HCC. Related to Figure 3A and Figure S3A.

**Table S6:** List of enriched Reacome, KEGG and Gene Ontology biological processes for deregulated pathways on the phosphoprotein levels. Related to Figure 3B and Figure S3B.

**Table S7:** List of kinases with altered activity in HCC and their associated substrates (at least a 5-fold difference between HCCs and normal livers) in a KSEA. Related to Figure 3C and Figure S3C.

**Table S8:** List of somatic mutations. Related to Figure 4 and Figure S4A.

**Table S9:** List of proteins differentially expressed between *CTNNB1*/*TP53*-mutant vs wild-type HCCs and their corresponding differential expression on the mRNA level. Related to Figure 4A/E.

**Table S10:** List of enriched Gene Ontology biological processes for *CTNNB1*- and *TP53*-mutant HCC. Related to Figure 4C/G.

**Table S11:** Pathway/KSEA analysis for single-omics subclasses. Related to Figure 5A-B and Figure S5H.

**Table S12:** Pathway/KSEA analysis for integrative clustering (iCluster) subclasses. Related to Figure 5C.

