## Supplemental information for "Proteogenomic characterization of hepatocellular carcinoma"

##### Figure S1

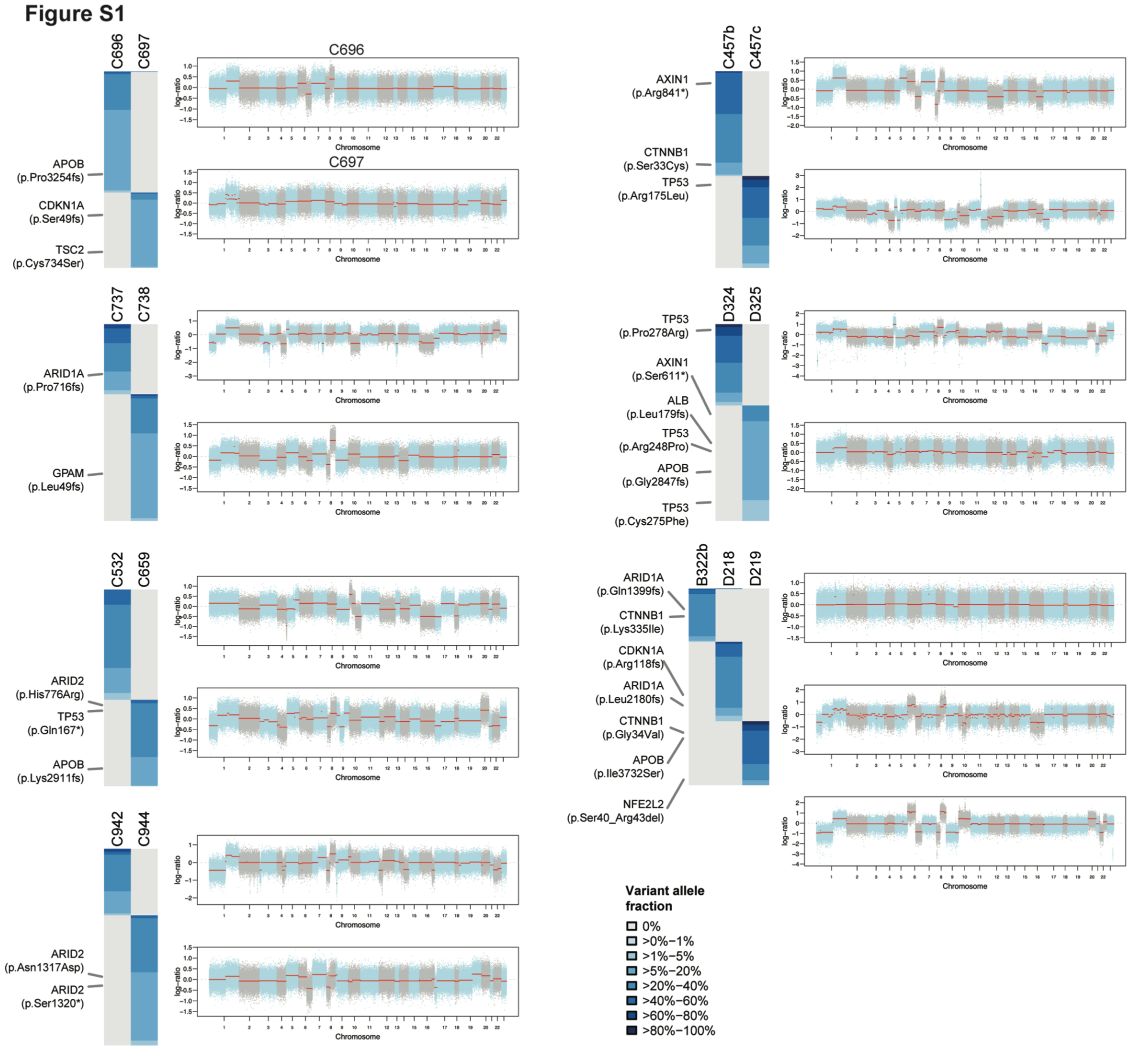


**Figure S1:** Heatmaps illustrate the variant allele fractions (shades of blue according to the color key, gray indicates absence) of the somatic mutations identified in the 7 patients for whom >1 tumor biopsy was included in the study. Non-synonymous mutations in HCC driver genes are labeled. To the right of the heatmaps are genome-wide copy number plots of the tumor biopsies. In the copy number plots, segmented Log2 ratios (y-axis) were plotted according to their genomic positions (x-axis). Alternating blue and gray demarcate the chromosomes. Related to Table 1.

### Figure S2

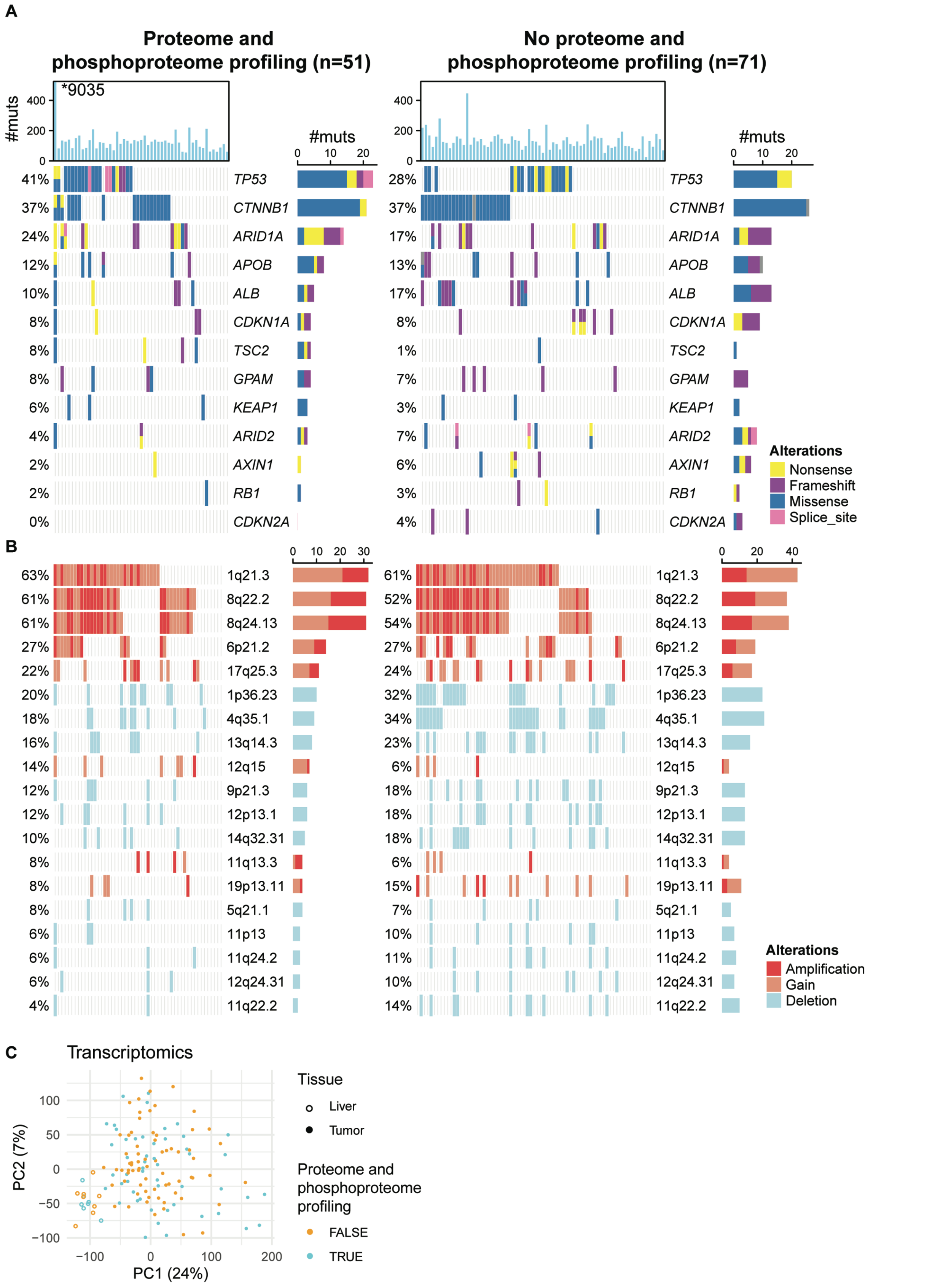


**Figure S2: (A)** Oncoprint showing the somatic mutational landscape of HCC biopsies, stratified by the availability of proteome and phosphoproteome profiling. Significantly mutated genes in the current cohort (*CTNNB1*, *CDKN1A*, *TP53*, *ALB*, *ARID1A*, *GPAM, AXIN1)* and six additional HCC driver genes (previously reported in at least 2 studies and mutated in at least 3 biopsies in this study) are included. Barplot above the oncoprint shows the total number of somatic mutations in each biopsy. Percentages to the left of the oncoprint show the fraction of biopsies harboring somatic mutations in a given gene. Barplot to the right of the oncoprint shows the total number and type of mutations identified in a given gene. The type of mutations is color-coded according to the legend. (**B**) Oncoprint showing the copy number profiles of HCC biopsies, stratified by the availability of proteome and phosphoproteome profiling. Significantly altered regions as defined by GISTIC2 on the entire cohort are shown. Copy number status was defined by GISTIC2. (**C**) Principal component analysis plot of gene expression of HCC biopsies (filled circles) and normal liver biopsies (open circles), colored by the availability of proteome and phosphoproteome profiling. Related to Table 1.

### Figure S3

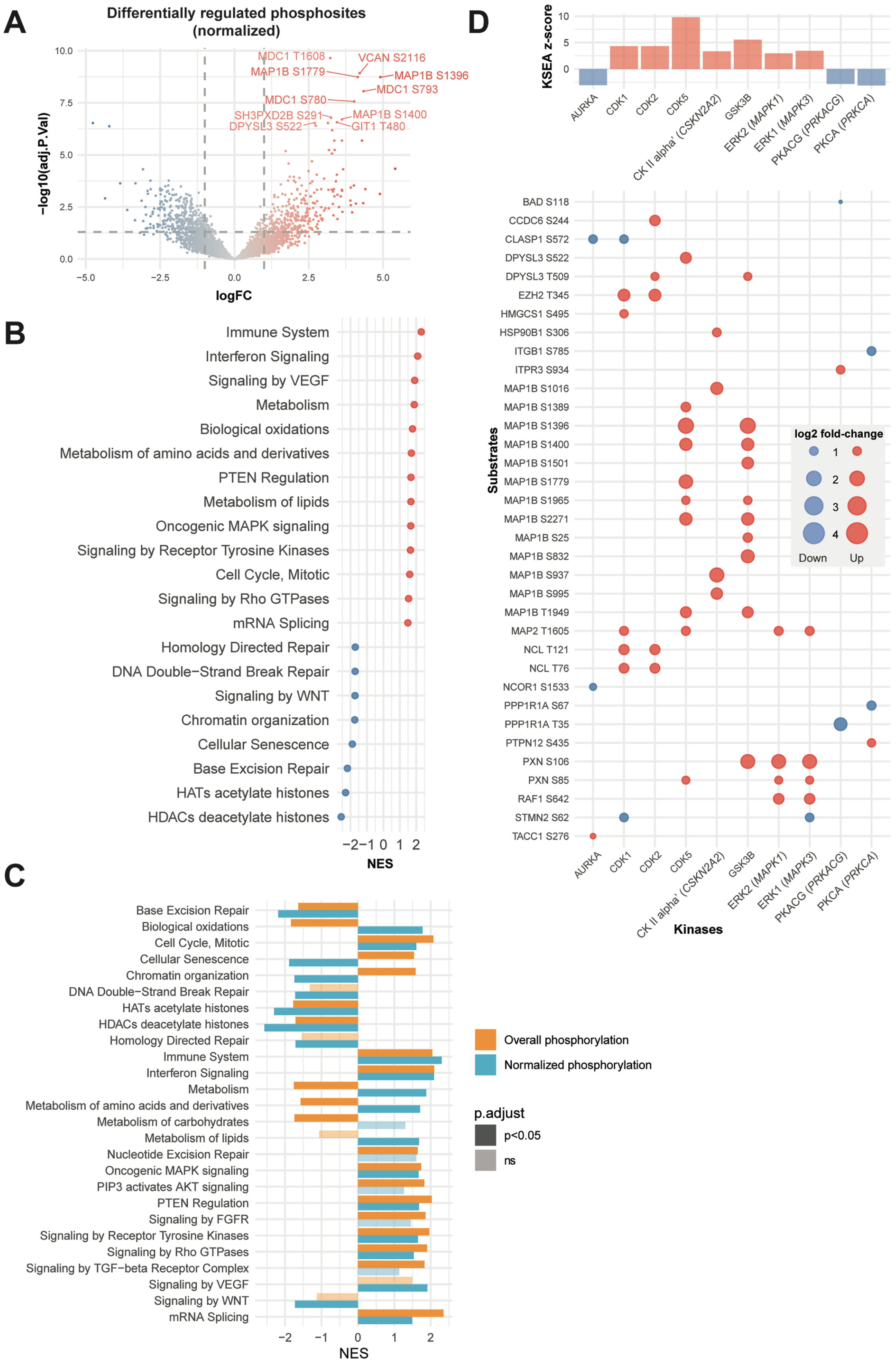
A
Differentially regulated phosphosites (normalized)
−log10(adj.P.Val)
logFC
MDC1 T1608
VCAN S2116
MAP1B S1779
MAP1B S1396
MDC1 S793
MDC1 S780
SH3PXD2B S291
MAP1B S1400
DPYSL3 S522
GIT1 T480
B
Immune System
Interferon Signaling
Signaling by VEGF
Metabolism
Biological oxidations
Metabolism of amino acids and derivatives
PTEN Regulation
Metabolism of lipids
Oncogenic MAPK signaling
Signaling by Receptor Tyrosine Kinases
Cell Cycle, Mitotic
Signaling by Rho GTPases
mRNA Splicing
Homology Directed Repair
DNA Double−Strand Break Repair
Signaling by WNT
Chromatin organization
Cellular Senescence
Base Excision Repair
HATs acetylate histones
HDACs deacetylate histones
NES
C
Base Excision Repair
Biological oxidations
Cell Cycle, Mitotic
Cellular Senescence
Chromatin organization
DNA Double−Strand Break Repair
HATs acetylate histones
HDACs deacetylate histones
Homology Directed Repair
Immune System
Interferon Signaling
Metabolism
Metabolism of amino acids and derivatives
Metabolism of carbohydrates
Metabolism of lipids
Nucleotide Excision Repair
Oncogenic MAPK signaling
PIP3 activates AKT signaling
PTEN Regulation
Signaling by FGFR
Signaling by Receptor Tyrosine Kinases
Signaling by Rho GTPases
Signaling by TGF−beta Receptor Complex
Signaling by VEGF
Signaling by WNT
mRNA Splicing
NES
Overall phosphorylation
Normalized phosphorylation
p.adjust
p<0.05
ns
D
KSEA z-score
AURKA
CDK1
CDK2
CDK5
CK II alpha' (CSKN2A2)
GSK3B
ERK2 (MAPK1)
ERK1 (MAPK3)
PKACG (PRKACG)
PKCA (PRKCA)
Substrates
BAD S118
CCDC6 S244
CLASP1 S572
DPYSL3 S522
DPYSL3 T509
EZH2 T345
HMGCS1 S495
HSP90B1 S306
ITGB1 S785
ITPR3 S934
MAP1B S1016
MAP1B S1389
MAP1B S1396
MAP1B S1400
MAP1B S1501
MAP1B S1779
MAP1B S1965
MAP1B S2271
MAP1B S25
MAP1B S832
MAP1B S937
MAP1B S995
MAP1B T1949
MAP2 T1605
NCL T121
NCL T76
NCOR1 S1533
PPP1R1A S67
PPP1R1A T35
PTPN12 S435
PXN S106
PXN S85
RAF1 S642
STMN2 S62
TACC1 S276
Kinases
log2 fold-change
Down
Up

**Figure S3: (A)** Volcano plot of the -log10(adjusted p-value) against the log fold-change (logFC) of the differentially regulated phosphosites normalized by overall protein levels ('normalized phosphosites') in HCC compared to normal livers. Dots are colored by logFC. Vertical dotted lines indicate |logFC|=2 and horizontal dotted lines indicate adjusted p-value=0.05. **(B)** Dot plot illustrating selected enriched Reactome pathways according to gene set enrichment analysis (GSEA) from the differential expression analysis in (**A**). NES: normalized enrichment score. **(C)** Top barplot showing the enrichment z-score of the kinases with significantly up- or downregulated kinase activity in a kinase-substrate enrichment analysis (KSEA) comparing normalized phosphosites in HCC to normal livers. In the bubble plot below, the phosphosite substrates are shown in rows, where red and blue dots indicate that the phosphosite is up- and downregulated, respectively. The size of the dots is proportional to the log2fold-change of the phosphosite. Phosphosites with at least a 5-fold difference between HCCs and normal livers are shown. For kinases with <3 substrates with at least a 5-fold difference, the top three substrates with the highest |logFC| are shown. Related to Figure 3.

#### Figure S4

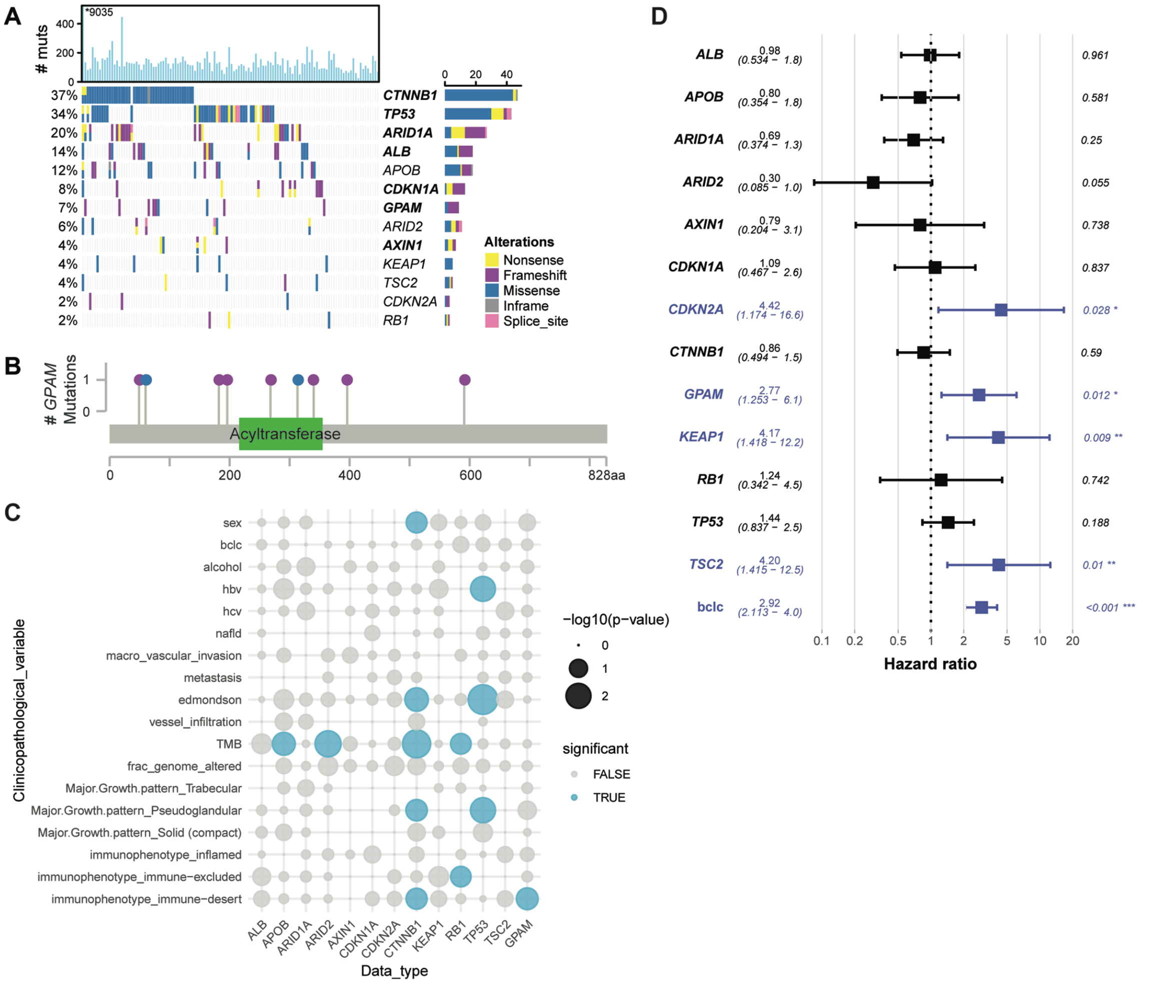


**Figure S4: (A)** Oncoprint showing the somatic mutational landscape of HCC. Significantly mutated genes in the current cohort (*CTNNB1*, *CDKN1A*, *TP53*, *ALB*, *ARID1A*, *GPAM, AXIN1*, in bold) and six additional HCC driver genes (previously reported in at least 2 studies and mutated in at least 3 biopsies in this study) are included. Barplot above the oncoprint shows the total number of somatic mutations in each biopsy. Percentages to the left of the oncoprint show the fraction of biopsies harboring somatic mutations in a given gene. Barplot to the right of the oncoprint shows the total number and type of mutations identified in a given gene. The type of mutations is color-coded according to the legend. **(B)** Lollipop plot showing the distribution of the *GPAM* mutations along the protein, with the mutations colored according to the color key in **(A)**. (**C**) Bubble plot showing association between mutation status and clinicopathological parameters. Size of the circles is proportional to -log10(p-value) and blue circles indicate statistically significant associations. Statistical analyses were performed by Fisher's exact or Chi-squared tests. (**D**) Forest plot showing multivariate Cox proportional-hazards model of overall survival according to the mutation status of HCC driver genes and BCLC clinical staging. Related to Figure 4.

**Figure S5**

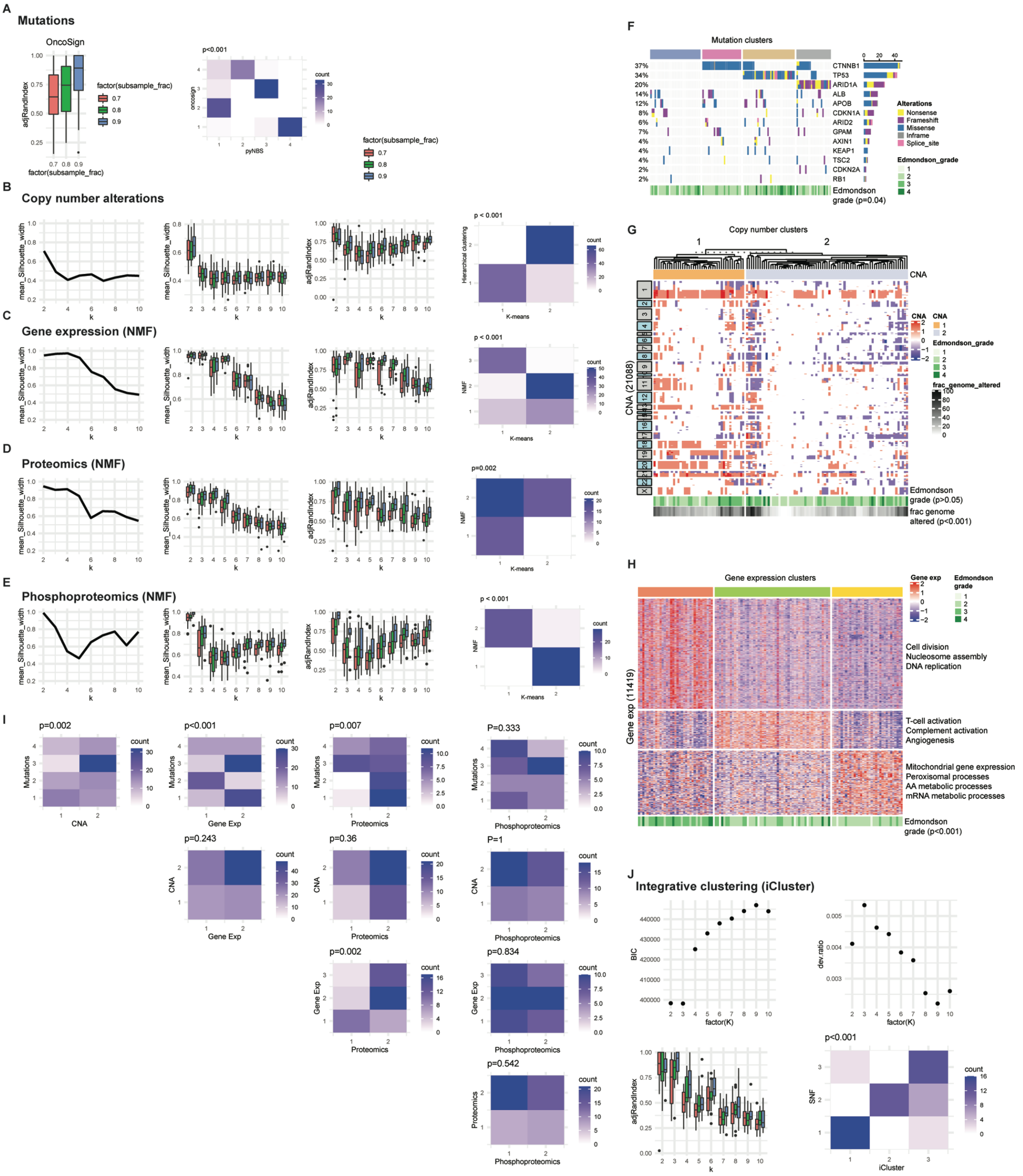


**Figure S5:** For the clustering of HCC biopsies based on somatic mutations, we used OncoSign (primary) and pyNBS (alternative). For CNA, we used consensus k-means clustering (primary) and consensus hierarchical clustering (alternative). For the remaining data types, we used consensus nonnegative matrix factorization (NMF, primary) and consensus k-means clustering (alternative). We assessed the mean Silhouette width (cluster quality) and adjusted Rand index (concordance with the classes derived from the full dataset) by subsampling 70%, 80% and 90% of the samples. (**A,** left) Boxplot showing adjusted Rand index between the classes derived from OncoSign (single-omics clustering using mutation data) using the full data set and the classes derived from downsampled data. Downsampling (70%, 80% and 90% of the samples) was performed over 20 iterations. (right) Concordance between clusters derived from OncoSign and pyNBS. (**B-E**, from left) Mean silhouette widths against k (the number of clusters), mean silhouette widths against k from clustering using downsampled datasets (70%, 80% and 90% of the samples, 20 iterations of downsampling), adjusted Rand index between the classes derived from the full dataset and the classes derived from subsampled data using the primary clustering method, and concordance between the primary and alternative clustering methods. Statistical comparisons were performed by Fisher's exact or chi-squared tests. **(F)** Clusters derived from OncoSign based on the significantly mutated genes found in this study and in previous studies (see methods). **(G)** Clusters derived from copy number alterations by consensus K-means clustering. **(H)** Clusters derived from gene expression by consensus NMF. **(I)** Concordance between single-omics clusters. Statistical comparisons were performed by Fisher's exact or chi-squared tests. **(J, top)** Assessment of the Bayesian Information Criterion and the deviance ratio (interpreted as percent explained variation) from integrative clustering using iClusterBayes. (bottom left) Boxplot showing adjusted Rank index between the classes derived from integrative clustering using the full data set and the classes derived from downsampled data. Downsampling (70%, 80% and 90% of the samples) was performed over 20 iterations. (bottom right) Concordance between clusters derived from iClusterBayes and SNF. Statistical comparison was performed using a chi-squared test. Related to Figure 5.

Figure S6

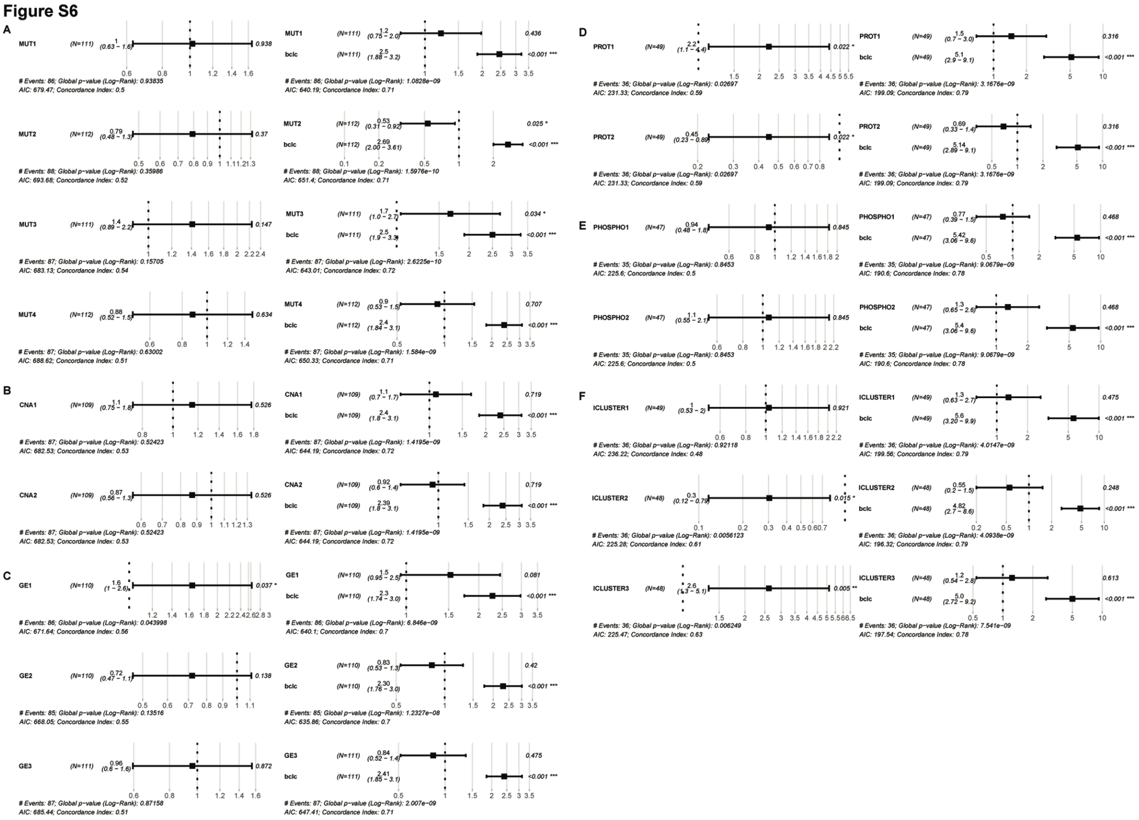

**Figure S6:** Forest plots from Cox proportional hazards analyses. (Left) Univariate analysis for each single-omics and integrative clusters. (Right) Multivariate analysis for each single-omics and integrative clusters incorporating BCLC clinical stage as a covariate. Related to Figure 5.
